# Transportability of mouse gene knockout models to human complex phenotypes

**DOI:** 10.64898/2026.11.04.764243

**Authors:** Laurence J Howe, Sara Wells, Neil M Davies, Sam M Lockhart, George Davey Smith

## Abstract

Mouse gene knockout studies are a valuable resource for understanding human gene function and the mechanisms underlying disease. However, translating insights to humans requires causal similarities between mouse and human phenotypes. A mouse model is transportable if gene perturbations that impact the mouse phenotype also affect (in orthologous genes) the corresponding human phenotype of interest, with consistent directionality.

To assess the transportability of mouse models to human complex traits we combined laboratory mouse gene perturbation data from the high-throughput International Mouse Phenotyping Consortium with predicted loss-of-function data from the UK Biobank within the Gene Perturbation Analysis for Transportability (GPAT) framework. Applying GPAT to 30 matched mouse-human phenotype pairs, we found evidence of transportability for 12 mouse phenotypes including biochemical (e.g. total cholesterol), cellular (e.g. lymphocyte counts) and anthropometric (e.g. length/height) phenotypes. In contrast, there was limited evidence for others such as fat mass and body mass index in the context of the murine experimental conditions. We also found some evidence of transportability of mouse glucose measurements to diabetes mellitus in humans.

Genome-wide perturbation effects on matched phenotypes were on average positively correlated between mice and humans (meta-analysis GPAT P = 2.3x10^-27^). However, the degree of correlation was generally modest and heterogeneous across phenotypes with some gene perturbations strongly associated in both species in opposite directions. Our framework illustrates how perturbation data can be used to quantitatively validate experimental models for understanding human disease aetiology and highlights the need for a range of orthogonal model systems, which may include challenges and perturbations to the mouse knockouts, to interrogate the mechanistic basis of human genetic associations.

## Main

Human genetics data on the impact of gene function changes (e.g., perturbations, knockouts) on disease risk and progression are a valuable resource for predicting therapeutic effects in drug discovery and development ^1–5^. Despite the emergence of large population biobanks with genome sequence data ^6,7^, understanding of the downstream effects of therapeutic (in)activation of the majority of protein-coding genes remains limited. Predicted loss-of-function (pLoF) variants for most genes are typically rare in the general population (frequency < 0.1%) and so effect estimates of gene function changes on human phenotypes from existing datasets are often highly imprecise particularly for uncommon or rare diseases ^5^.

Mice are a useful model organism for understanding effects of human gene function changes because of the substantial orthology between the mouse and human genomes, with more than two thirds of human protein coding genes having a corresponding high-confidence orthologue in mice. Experimental capabilities in mice enable direct estimates of the impact of gene knockouts on mouse phenotypes with extremely precise environmental control, on a homogenous genetic background ^8–15^, potentially helping to overcome difficulties in proving causality in humans. Mouse phenotypic screen data are commonly integrated into clinical pipelines with evidence of value in improving diagnostic yield of human Mendelian diseases ^8^. Mouse knockout data can also provide biological insights into putative targets for human complex diseases. However, in order for mouse models to be informative for human therapeutics development, the effects of perturbing mouse genes need to be consistent with effects on human phenotypes ^8,14,16^. Indeed, a complex phenotype such as adiposity may have distinct genetic aetiology in humans and mice because of environmental factors (e.g., experimental conditions) or genuine interspecies biological differences in gene function.

The transportability of a mouse gene-knockout model to a human phenotype is the extent to which gene perturbation effects on phenotypes in mice are consistent with effects of (orthologous) gene perturbations on the corresponding human phenotype ^17^. For example, mouse gene knockout effect estimates on total cholesterol are transportable to humans if gene perturbations impacting total cholesterol in mice also affect (in orthologous genes) total cholesterol in humans with consistent directionality. If a model is transportable, then mouse gene perturbation data is an informative predictor of the effects of therapeutic (in)activation of a gene on the human phenotype.

There has been limited work evaluating the extent to which inbred mouse knockout models are transportable to human phenotypes using gene perturbation data. If we can better understand when mouse models are transportable, we can better incorporate mouse data into developing and prioritising novel therapeutics and treatments or modify the conditions and challenges that mice are exposed to in order to better recapitulate the translational human context. We apply the Gene

Perturbation Analysis for Transportability (GPAT) framework ^17^, a gene-level implementation of Mendelian randomization ^18,19^, to assess transportability of mouse models to corresponding matched human phenotypes. We also evaluate the transportability of diabetes-related mouse models to diabetes and biomarkers in humans. We use estimates of gene perturbation effects from the International Mouse Phenotyping Consortium (IMPC) dataset ^11^ and from UK Biobank Whole-Genome Sequence (WGS) ^20,21^ burden tests (humans).

### Mouse and human perturbation data

The IMPC database ^11^ (v-24.0) is a high-throughput dataset from a large number of strains carrying null alleles for single genes on a standardised inbred background (C57BL/6N). The data is collected from mouse research centres globally, performing controlled phenotyping pipelines producing data across multiple body systems. The phenotypes are limited by age (most pipelines end at 16 weeks of age) and the mice are housed in standard laboratory specific pathogen-free conditions. Control wild type mice and phenotyped periodically, delivering a rolling baseline and statistical analysis performed using a windowing approach that assigns higher weights to control mice measured close in time to the mutant mice ^22^. The dataset contains gene perturbation data (heterozygous and homozygous variants) for 1,827,009 gene-pair phenotype pairs covering 9881 mouse genes and > 1500 mouse phenotypes. Of these, there is evidence that 3970 unique genes affect 566 phenotypes (at a threshold of P < 1x 10^-5^). UK Biobank ^20^ is a population study of around 500,000 individuals with a wide range of measured phenotypes and WGS data. UK Biobank WGS data provides rare pLoF evidence for >1000 gene-phenotype pairs ^7^. The dataset includes both heterozygous and homozygous carriers although homozygous pLoF carriers are rare for most genes.

We manually reviewed the mouse phenotypes ^23^ with perturbation evidence to pair them with human phenotypes measured in UK Biobank. We manually matched 30 phenotypes with 28 having the same definitions in mice and humans (e.g. total cholesterol). The two exceptions were ‘body length, MP:0001257/1258’ in mice being paired with standing height in UK Biobank (data showcase ID: 50) and ‘body weight, MP:0001260/1262’ paired with BMI (data showcase ID: 21001). Unpaired phenotypes which were not analysed included mouse behavioural phenotypes which are not straightforward to match with human phenotypes (e.g. abnormal tail dragging, MP:0005507) or phenotypes which were not measured in UK Biobank participants (e.g. circulating magnesium levels, MP:0010092/10093) (**Supplementary Table 1**).

Mouse and human genes were matched based on high confidence orthology and one-to-one mappings (**Methods**). Overall human genes with orthologues in the IMPC database had a similar cumulative pLoF frequency to genes not in the IMPC data (P = 0.20). However, human genes with orthologues in the IMPC database with perturbation evidence for one or more mouse phenotypes (3483 genes, P < 1x10^-5^) had a lower cumulative pLoF frequency (ratio of mean frequencies = 0.85, P = 6.1x10^-4^). These results suggest mouse models can enable insights into constrained human genes with limited human pLoF data.

### Transportability of paired mouse models

We applied the GPAT framework ^17^ to evaluates transportability between mouse models and paired human phenotypes. GPAT is a gene-level implementation of Mendelian randomization (MR) evaluating the consistency between gene perturbation effects on a mouse phenotype and gene ortholog perturbation effects on the paired human phenotype.

Assume there are *j ɛ J* genes where perturbations are strongly associated with a mouse phenotype and we have harmonised gene perturbation effect estimates on the mouse (*β_j,k_*) and human (*β_j,/_*) phenotypes. Wald ratios for each gene, which are ratios of the effect sizes, have coefficients 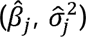:

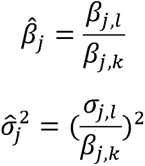

Mouse gene knockout effect estimates are often collected from different studies. We observed that effect estimates for the same phenotype in different strains were highly heterogeneous, due to differences in experimental details such as experimental protocol, units of measurement, housing, husbandry, equipment differences and covariates. Outlying effect sizes can lead to extreme Wald ratios. However, there was insufficient data to control for this heterogeneity at scale so we used unweighted directional coefficients 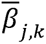 to compute Wald ratios instead of the raw weighted effect sizes *β_j,k_* (**Methods**). This means that the approach is modelling only directionality but not quantitative effect sizes from the mouse data with both directionality and quantitative effect sizes modelled from the human data.

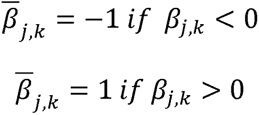

GPAT estimates are combinations of Wald ratios across all genes. For example, using a fixed-effects inverse-variance weighted meta-analysis where the coefficients 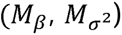 are a combined estimate of equivalence between mouse and human perturbation effects across all J genes:

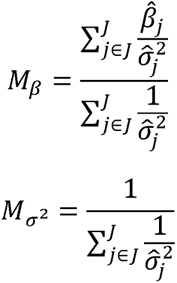

After data harmonisation steps (**Methods**), we performed GPAT analyses for 30 phenotype matched mouse-human pairs. Fixed-effects GPAT estimates provided evidence of transportability (GPAT fixed-effects P < 0.05, GPAT beta > 0) of mouse models to humans for 12 phenotypes; total cholesterol, triglycerides, bilirubin, alkaline phosphatase, lymphocyte counts, mean corpuscular volume, platelet counts, mean platelet volume, length-height, albumin, red blood cell width and bone mineral density (**Figure 1A / Supplementary Table 2**). Across all 30 phenotypes, a meta-analysis of the fixed-effects GPAT estimates indicated that on average mouse estimates are positively correlated with human pLoF estimates for the tested phenotype set (beta = 0.009, P-value = 2.3x10^-27^). There was limited GPAT evidence to support transportability of mouse models under the experimental conditions for a number of phenotypes despite a large number (>50) of mouse knockout genes associated with each phenotype. For example, BMI, fat-mass, HDL-cholesterol and glucose.

**Figure 1A.**
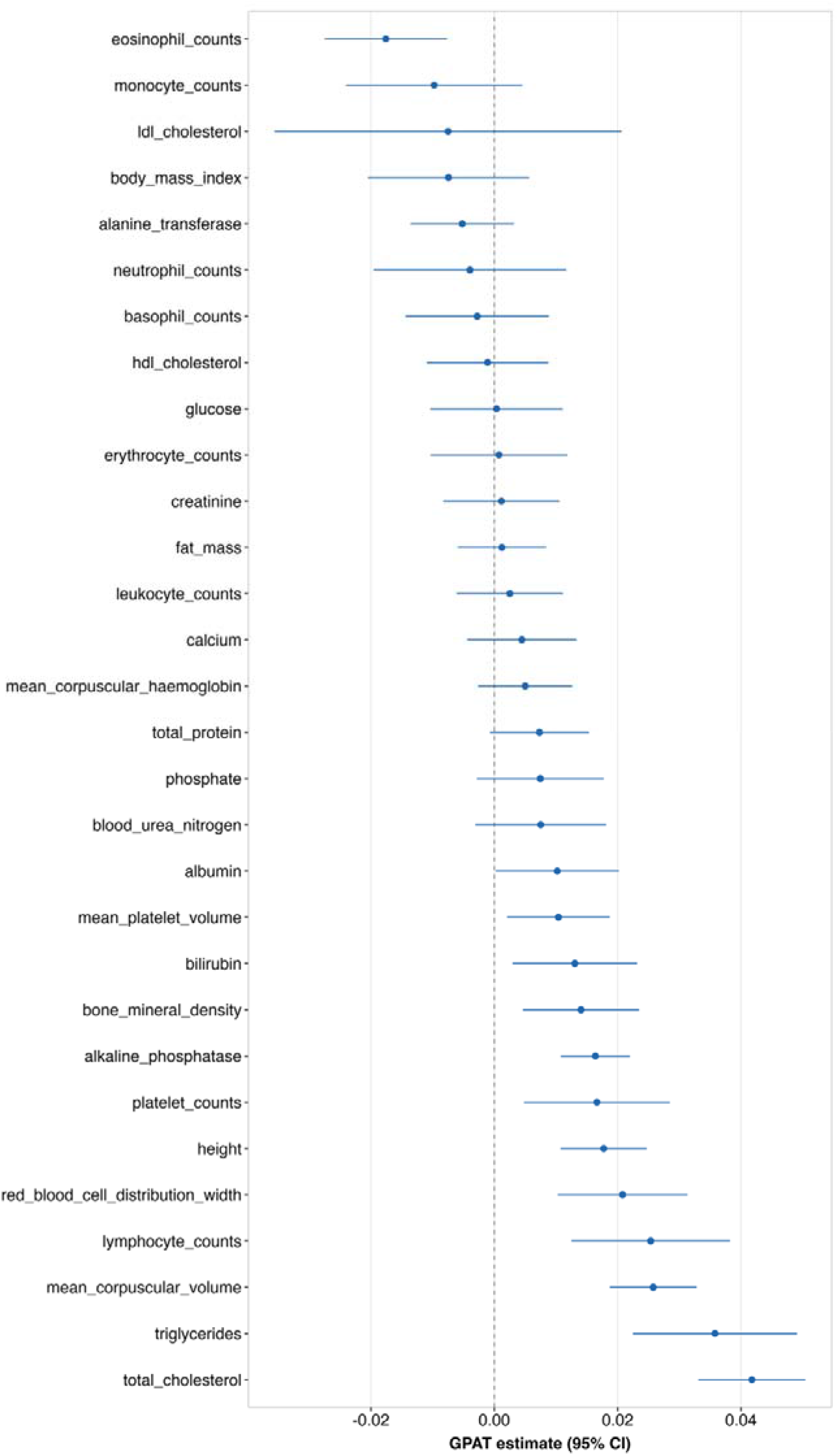
Fixed-effects GPAT estimates for 30 mouse-human phenotype pairs Figure 1A presents fixed-effects GPAT estimates and 95% confidence intervals for 30 pairs of matched mouse and human phenotypes. GPAT estimates capture (dis)similarities between gene-phenotype effect estimates. The Y axis contains human phenotype names. The presented alkaline phosphatase estimate excludes *GPLD1* (see discussion in text). These estimates model only direction of effect in mice and both direction of effect and magnitude of effect in humans.

**Figure 1B.**
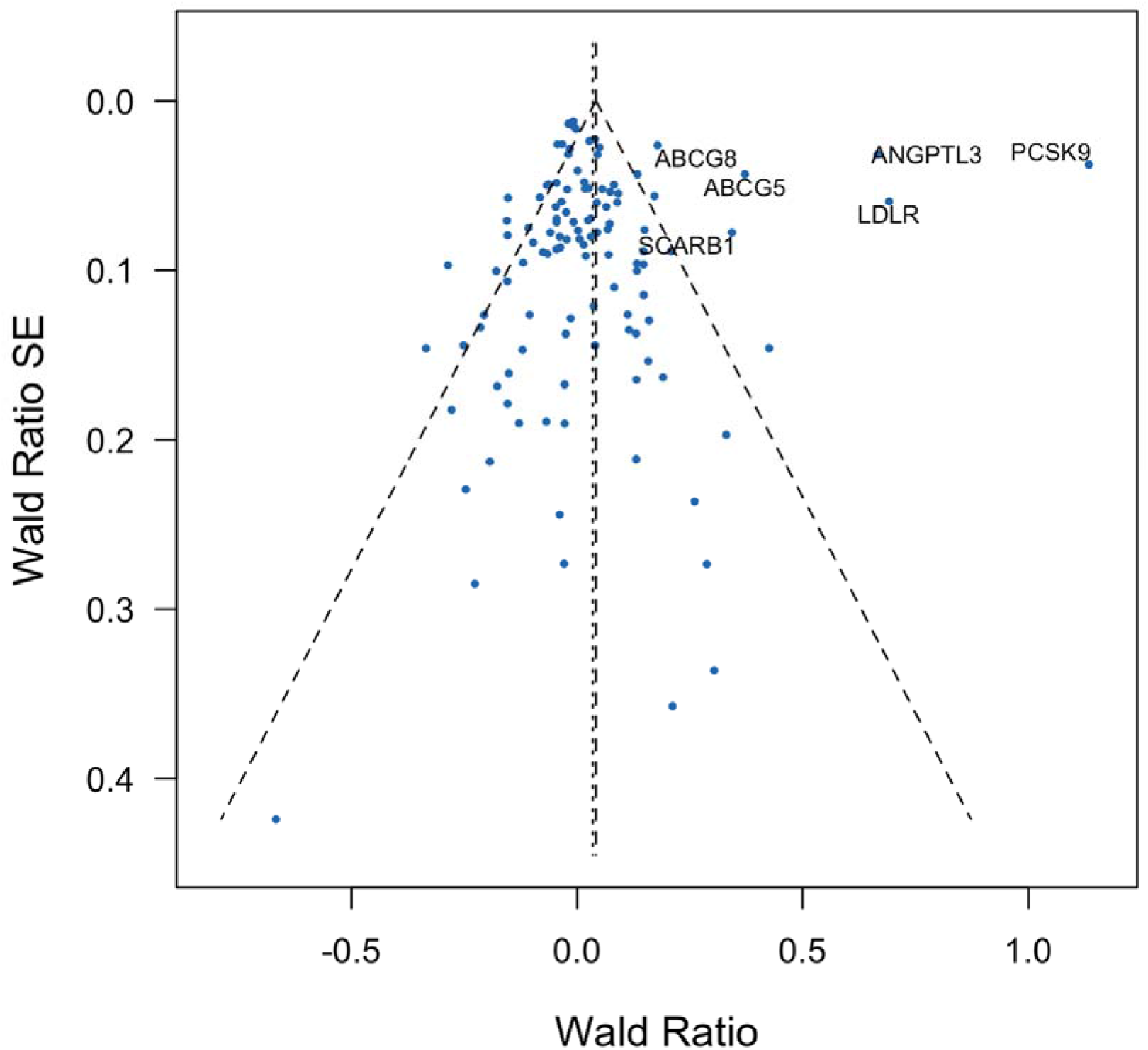
Gene perturbations strongly associated with total cholesterol in mice and their association with total cholesterol in humans Figure 1B presents Wald ratios and standard errors for individual genes strongly associated with total cholesterol in mice. Wald ratios are the gene-phenotype effect estimate in humans divided by the gene-phenotype effect estimate in mice. We used directional, unweighted mouse estimates and so the Wald ratios are the effect estimates from humans with positive (consistent directionality between mice and humans) or negative (inconsistent directionality) values. The dotted vertical lines indicate the overall fixed and random effects GPAT estimates across all genes. The dotted diagonal lines represent 95% confidence intervals. All genes with pLoF burden P < 0.001 in humans are annotated with the human gene symbol. These estimates model only direction of effect in mice and both direction of effect and magnitude of effect in humans.

**Figure 1C.**
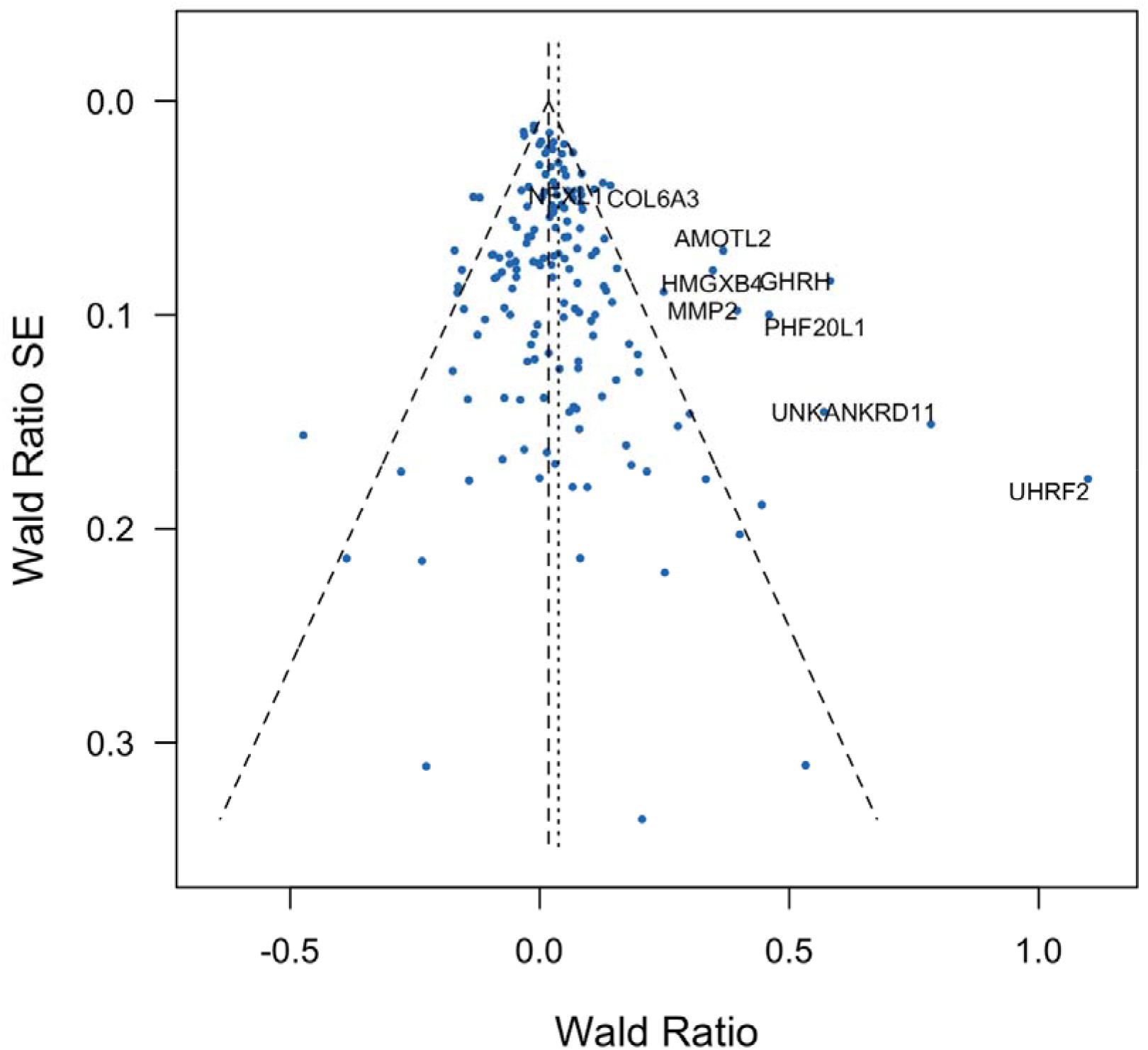
Gene perturbations strongly associated with length in mice and their association with height in humans Figure 1C presents Wald ratios and standard errors for individual genes strongly associated with length in mice. Wald ratios are the gene-phenotype effect estimate in humans divided by the gene-phenotype effect estimate in mice. We used directional, unweighted mouse estimates and so the Wald ratios are the effect estimates from humans with positive (consistent directionality between mice and humans) or negative (inconsistent directionality) values. The dotted vertical lines indicate the overall fixed and random effects GPAT estimates across all genes. The dotted diagonal lines represent 95% confidence intervals. All genes with pLoF burden P < 0.001 in humans are annotated with the human gene symbol. These estimates model only direction of effect in mice and both direction of effect and magnitude of effect in humans.

**Figure 1D.**
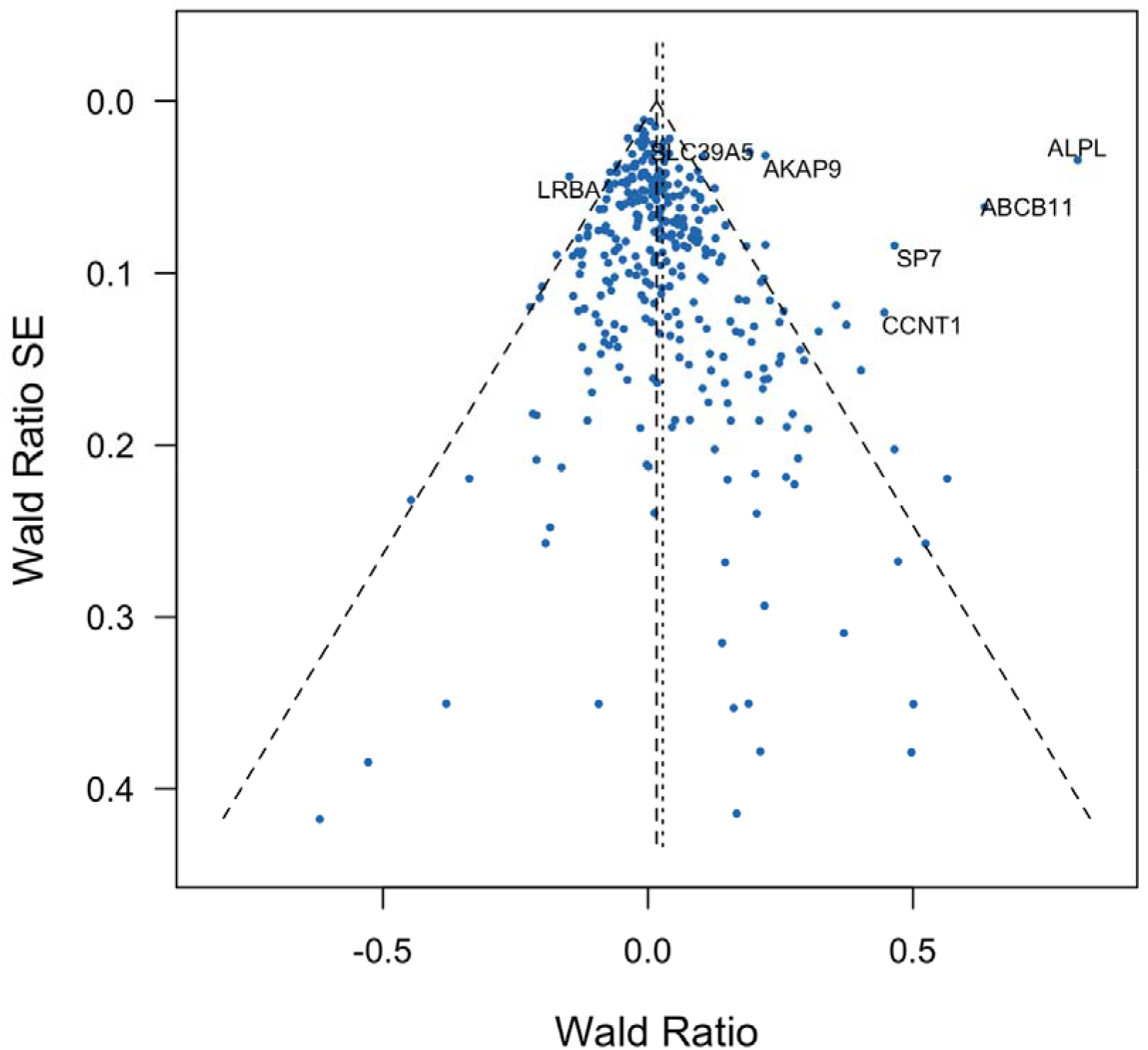
Gene perturbations strongly associated with alkaline phosphatase in mice and their association with alkaline phosphatase in humans Figure 1D presents Wald ratios and standard errors for individual genes strongly associated with alkaline phosphatase in mice. Wald ratios are the gene-phenotype effect estimate in humans divided by the gene-phenotype effect estimate in mice. We used directional, unweighted mouse estimates and so the Wald ratios are the effect estimates from humans with positive (consistent directionality between mice and humans) or negative (inconsistent directionality) values. The dotted vertical lines indicate the overall fixed and random effects GPAT estimates across all genes. The dotted diagonal lines represent 95% confidence intervals. This funnel plot excludes *GPLD1* which was removed in the primary analysis but has an effect in the complete opposite direction. All genes with pLoF burden P < 0.001 in humans are annotated with the human gene symbol. These estimates model only direction of effect in mice and both direction of effect and magnitude of effect in humans.

Phenotypes where GPAT evidence indicated transportability of mouse models featured multiple gene perturbations which influenced the phenotype in both mice and humans with consistent direction of effect. However, these phenotypes also featured gene perturbations which were strongly associated in mice but had effects in the opposite direction in humans or very limited evidence of an association despite sufficient statistical power. For example, for total cholesterol, perturbation of well-characterised lipid genes (*ANGPTL3, LDLR, PCSK9*) had consistent effects between mice and humans and resulted in a positive GPAT estimate. However, there were perturbations which had inconsistent interspecies effects such as *NR1H4* which was strongly associated with increased circulating cholesterol in mice (P < 1x10^-50^) but had some evidence of decreasing total cholesterol in humans (beta = -0.39; P = 0.0035). In addition, there were many gene perturbations with strong effects on mice’s cholesterol which had little effect in humans despite abundant pLoF carriers (**Figure 1B**). Gene perturbations influencing length in mice generally had consistent effects on height in humans but again consistency across genes was heterogeneous (**Figure 1C**).

Alkaline phosphatase had a raw negative fixed-effects GPAT estimate (beta = -0.018; P = 2.3x10^-10^) while the random-effects estimate was in the opposite direction (beta = 0.026; P = 0.0010). The discordance was a consequence of discordance in directionality for *GPLD1*, a phospholipase targeting GPI-anchored proteins. Loss-of- function is strongly associated with higher levels of alkaline phosphatase in mice but extremely strongly associated with lower levels in humans (P < 1x10^-200^). Several genes strongly associated in humans, including *ALPL* (Alkaline Phosphatase gene), had concordant effect directions between mice and humans. For the purposes of the at-scale fixed-effects reporting we excluded *GPLD1* due to its extreme heterogeneous (relative to other genes) impact on the overall GPAT estimate and presented the resulting fixed-effects GPAT estimate (beta = 0.016; P = 9.9x10^-9^) (**Figure 1D**). We further explore the prevalence of directional concordance later in the manuscript.

GPAT fixed-effects estimates indicated a negative correlation between mice and human perturbation effects for eosinophil counts (beta = -0.018, P = 5.1x10^-4^). This negative estimate was driven by *RASAL3* where perturbation was strongly associated with higher eosinophil differential count in mice but associated with lower eosinophil counts in humans (*RASAL3* beta = -0.08, P = 4.8x10^-9^). Reviewing the IMPC mouse data, the increasing effect for *RASAL3* was observed in females (P = 5.2x10^-7^) but not males (P = 0.64). The female estimate appeared to be driven by 2 (out of 8) mutants with extremely high eosinophil counts and the effect estimate in males was in the opposite direction suggesting outlier issues although we cannot rule out partial gene penetrance (**Supplementary Figure 1**).

We next performed cross-phenotype GPAT analyses to evaluate the transportability of total cholesterol mouse gene knockout data to the set of 30 human phenotypes. We found strong evidence of transportability of total cholesterol in mice to LDL-cholesterol (GPAT beta = 0.045, P = 5.7x10^-25^) and triglycerides (GPAT beta = 0.025, P = 4.6x10^-8^) in humans. This contrasts with the weak evidence (**Figure 1**) of transportability of mouse LDL-cholesterol to human LDL-cholesterol. These results likely reflect both the shared aetiology between lipid measures and the more comprehensive gene coverage in the knockout database for total cholesterol (414 genes for total-cholesterol versus 72 for LDL-cholesterol). Indeed, key lipid genes such as *LDLR* and *PCSK9* lacked mouse knockout data specifically for LDL-cholesterol but did for total cholesterol.

There was also some evidence (GPAT P-value < 0.05) of transportability of total cholesterol mouse genes to adiposity phenotypes (BMI, fat mass), red blood cell distribution width and alkaline phosphatase. The adiposity evidence is striking, as GPAT with matched mouse phenotypes (i.e. BMI-BMI and fat mass – fat mass) (**Figure 1A**) provided no evidence of transportability (**Figure 2** / **Supplementary Table 3**).

**Figure 2.**
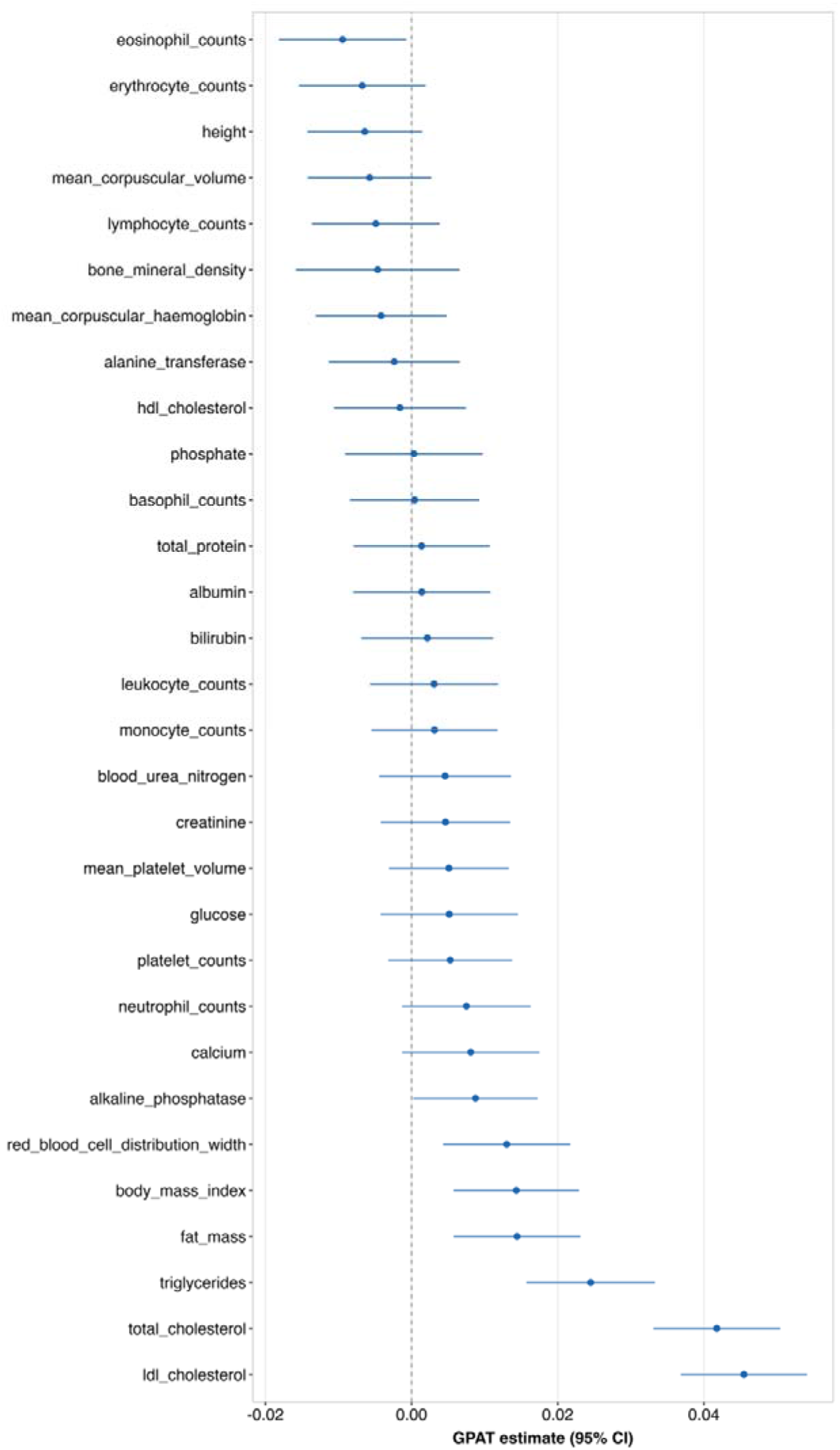
Fixed-effects cross-phenotype GPAT estimates for total cholesterol in mice on human phenotypes Figure 2 presents fixed-effects GPAT estimates and 95% confidence intervals for mouse total cholesterol with 30 human phenotypes. GPAT estimates capture (dis)similarities between gene-phenotype effect estimates. The Y axis contains human phenotype names. These estimates model only direction of effect in mice and both direction of effect and magnitude of effect in humans.

To investigate cross-phenotype associations within mice, we also evaluated whether total cholesterol knockouts were associated with other murine phenotypes with consistent directionality. The mouse effect size estimates are heterogeneous and there is sparse coverage of gene-phenotype data within the database. Therefore, we focused on concordance in sign without modelling effect sizes. We used a binomial sign test on 131 genes for total cholesterol (P < 1x10^-5^) with available data for each of the 29 murine phenotypes (N between 4 and 102). There was evidence (P < 0.05) of consistent directions of effect for total cholesterol with subfractions (HDL cholesterol, LDL cholesterol) as well as for total protein, triglycerides, calcium, albumin and glucose. Stronger conclusions for some phenotypes were limited by available data. For example, for body weight and basophils, only 4 and 5 of the total cholesterol genes had available data (**Supplementary Table 4**).

### Directional consistency of perturbation effects between mice and humans

GPAT estimates provide insight into transportability of mouse models by combining evidence across a set of gene perturbations strongly associated with a mouse phenotype. GPAT fixed-effects estimates can be strongly impacted by perturbations which are associated in both mice and humans but in opposite directions. This can lead to misleading fixed-effects estimates if a gene has an extremely large test statistic in humans such as *GLPD1* and alkaline phosphatase.

There were 323 perturbation-phenotype pairs where the perturbation was strongly associated in mice (P < 1x10^-5^) and nominally associated in humans with the matched phenotype (pLoF burden P < 0.05) (**Supplementary Table 5**). In this set, 214 (66.3%) had effects in the same direction indicating general directional concordance but also substantial heterogeneity. Restricting this sample to 45 pairs with stronger pLoF burden evidence in humans (P < 1x10^-5^), 38 (84%) of the pairs were directionally concordant with the 7 highly discordant pairs including the two examples discussed above (*GPLD1*, *RASAL3*), three perturbations associated with mean platelet volume (*BIN2, FBXO38, SLFN14*) and two perturbations associated with mean corpuscular haemoglobin (*RHAG, SPTB*). Manual review of the individual-level data scatter plots for some of these gene-phenotype pairs (*RASAL3, FBXO38, SLFN14*) suggested that the strong mouse associations may be driven by outliers. However, it is difficult to distinguish outliers from partial gene penetrance where only a subset of individuals display the phenotype.

We empirically investigated two possible explanations for directional inconsistency, pLoF mis annotation in humans and orthology differences across matched mouse-human gene pairs.

To assess possible pLoF mis annotation in humans, we evaluated the association between gene pLoF and plasma protein-levels using UK Biobank data (N ∼ 40K) on 2120 proteins where the relevant gene had 1 or more LoF carriers. Considering the 109 genes with discordant associations (human P < 0.05), 12 of these had pLoF burden data for the encoded plasma protein levels. Burden estimates were generally consistent with pLoF lowering plasma protein-levels (median Z score: -4.6; Q1-Q3: - 7.3, -1.9) with no strong evidence that pLoF increases plasma protein-levels for any of the tested genes. These results suggest that the observed interspecies discordance in perturbation effects is unlikely to relate to mis annotation of pLoFs in humans.

For matched mouse-human gene pairs, we restricted to high-confidence orthologs with one-to-one matchings but within this set there is variation in the percentage of identical sequence between the mouse and human genomes. Comparing the sets of 214 concordant and 109 discordant genes we observed slightly lower mouse orthology % for the discordant pairs (concordant mean: 86.5%, discordant mean: 84.8%, P = 0.14) limiting stronger inference. These results suggest that minor differences in mouse and human genomes are unlikely to explain all of the observed discordance in gene-phenotype associations.

### Transportability of mouse models to diabetes mellitus in humans

Mouse knockout studies are often used to evaluate perturbation effects on endophenotypes measurable in mice to better understand the aetiology of human diseases. For example, mouse glucose phenotypes and type II diabetes. GPAT analyses can be used to evaluate cross-trait transportability of mouse phenotypes to human diseases providing more direct evidence for value of mouse knockout data in disease-focused drug development.

We investigated transportability of mouse models to diabetes mellitus in humans. For diabetes-related mouse phenotypes we considered gene-knockouts associated with circulating free-fed glucose, fasting glucose (measured at time zero in an Intraperitoneal glucose tolerance test: IPGTT), glucose tolerance (area under curve in IPGTT; higher values correspond to lower glucose tolerance) and circulating blood insulin. We divided these genes into five sets; glucose all (N = 295): genes associated with free-fed glucose, fasting glucose and/or glucose tolerance; free-fed glucose (N = 119); fasting glucose (N = 144); glucose tolerance (N = 94); and insulin (N = 14). In humans we considered a binary diabetes outcome (any diabetes including type I and type II) and two continuous biomarkers; Hba1c a continuous diagnostic for type II diabetes, and the ratio of Triglycerides to HDL-cholesterol (TG:HDL), a marker of insulin resistance which is readily available in population biobanks ^24^.

There was some evidence of transportability of the glucose tolerance gene-set to diabetes (GPAT beta = 0.055; P = 0.042). In contrast, evidence of transportability to diabetes for the glucose all, free-fed glucose, fasting glucose and insulin gene-sets was more limited. The fasting glucose (GPAT beta = -0.010; P = 0.016) gene set was associated with Hba1c but in the opposite direction to expected and the glucose all and free-fed glucose gene sets were associated in the opposite direction with TG:HDL ratio (GPAT betas = -0.009 & -0.013; P = 0.006, 0.010). The insulin gene set was not strongly associated with any of the human phenotypes but limited by the modest number of genes with mouse knockout evidence in the database (**Figure 3; Supplementary Figures 2-15; Supplementary Table 6**).

**Figure 3.**
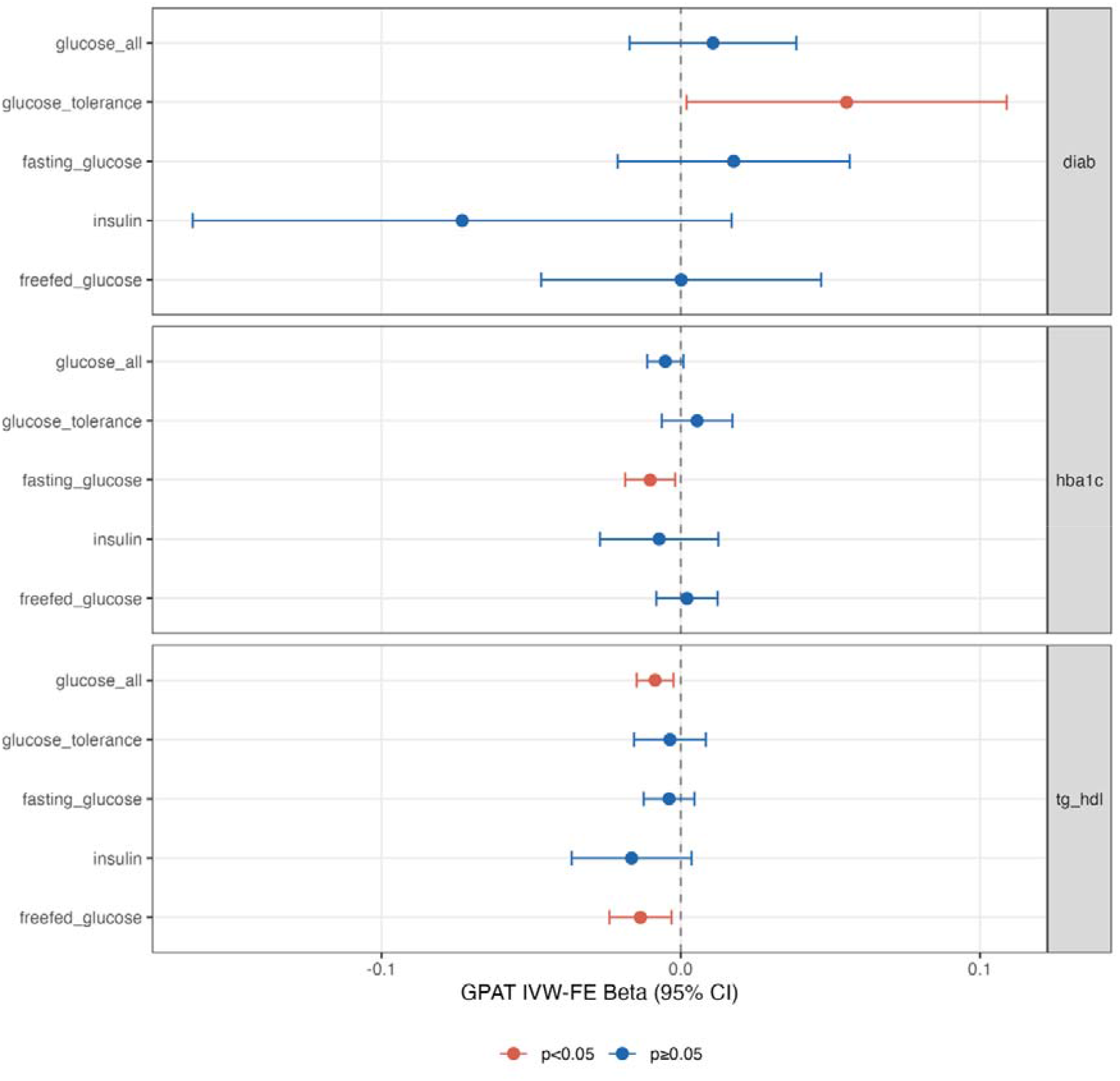
Gene perturbations strongly associated with glucose and insulin phenotypes in mice and their association with diabetes-related phenotypes in humans. Figure 3 presents fixed-effects GPAT estimates and 95% confidence intervals for mouse glucose and insulin-related genes with diabetes mellitus (diab), Hba1c (hba1c) and TG:HDL (tg_hdl). GPAT estimates capture (dis)similarities between gene-phenotype effect estimates. These estimates model only direction of effect in mice and both direction of effect and magnitude of effect in humans.

These results suggests that dynamic glucose tolerance tests in mice may be a more translatable phenotype than simpler assessments of fasting glucose and HbA1c. Two genes driving the evidence for transportability of mouse phenotypes to diabetes were *GCK: Glucokinase* (human diabetes beta = 1.98; P = 1.5x10^-14^) and *RFX6* (human diabetes beta = 1.13; P = 5.6x10^-4^), both established MODY genes ^25^ with perturbation of both associated with impaired glucose tolerance in mice and increased risk of diabetes in humans (**Figure 4**). Notably burden tests of protective effects on disease are often underpowered so we are likely insufficiently powered to replicate (in humans) some putative protective effects observed in mice.

**Figure 4.**
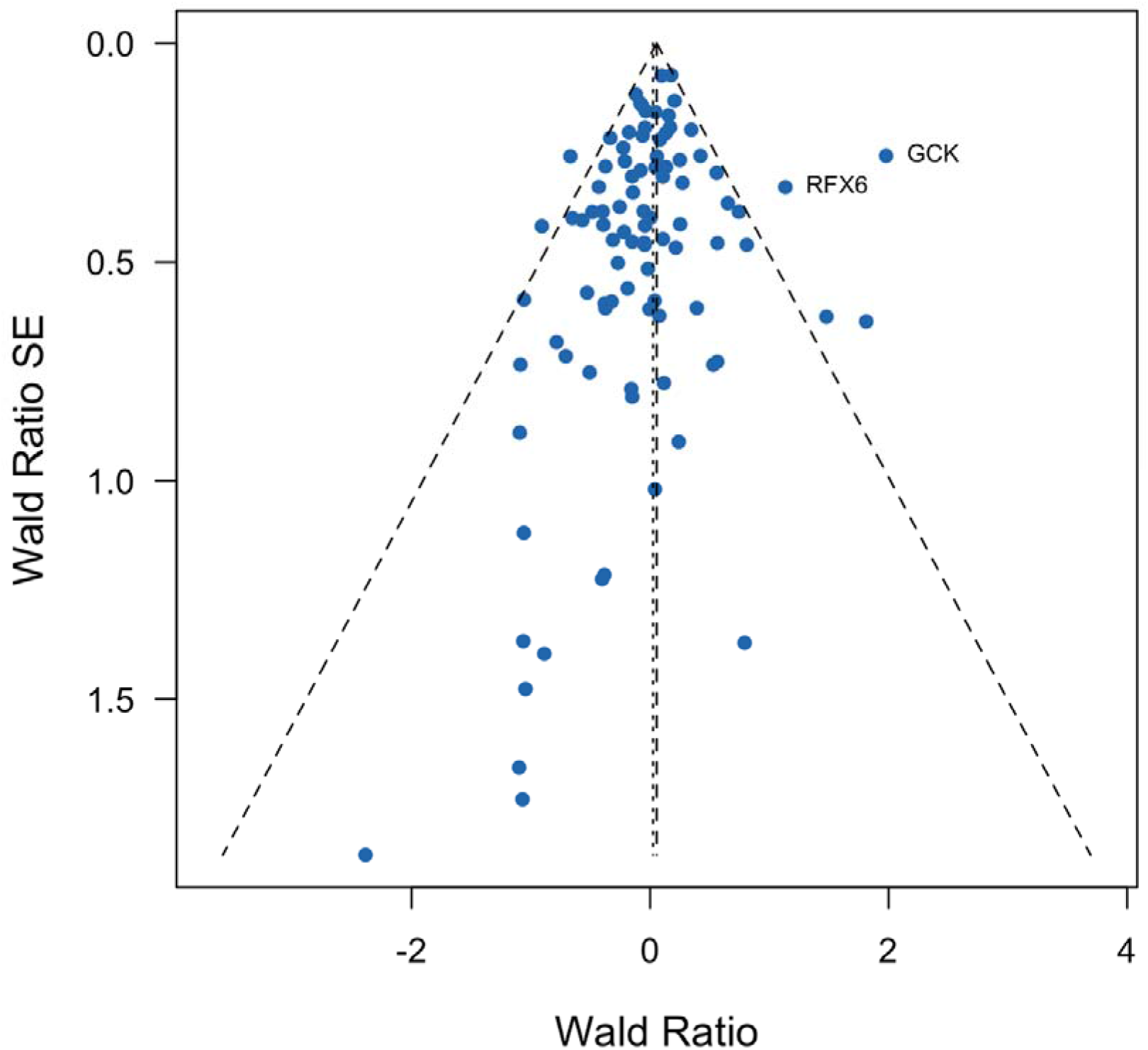
Gene perturbations strongly associated with glucose tolerance in mice and their association with diabetes mellitus in humans Figure 4 presents Wald ratios and standard errors for individual genes strongly associated with glucose tolerance in mice with respect to diabetes mellitus in humans. Wald ratios are the gene-phenotype effect estimate in humans divided by the gene-phenotype effect estimate in mice. We used directional, unweighted mouse estimates and so the Wald ratios are the effect estimates from humans with positive (consistent directionality between mice and humans) or negative (inconsistent directionality) values. The dotted vertical lines indicate the overall fixed and random effects GPAT estimates across all genes. All genes with pLoF burden P < 0.001 in humans are annotated with the human gene symbol.

## Discussion

A motivating aim of model organism research is to better understand the aetiology of human phenotypes informing effects of interventions ^8,13^. For example, there is growing interest in using *in-vivo* gene perturbation estimates from non-human animals to inform the effects of the orthologous gene in humans. A previous study performed in-vivo CRISPR base editing of *PCSK9* in primates before commencing clinical trials in humans ^26^. However, such applications require assuming that the non-human model is transportable to the human phenotype of interest. Previous research has used semantic similarity-based approaches on free-text to assess similarities between mouse and human phenotypes ^27–29^. Here we propose an orthogonal approach to quantitatively test the transportability assumption using gene perturbation data. If a model is transportable then we can potentially extrapolate findings to humans. We have partial data on how gene perturbations in humans affect a wide range of phenotypes which can be used as a pseudo-benchmark dataset. Data are underpowered for many individual gene-phenotype pairs but by combining data across many genes we can assess how well perturbation data for a non-human phenotype model corresponds to observed effects on that phenotype in humans.

Using perturbation data we applied the GPAT framework ^17^ to evaluate model transportability of mouse models from a large gene-knockout database to corresponding matched human phenotypes. We found evidence of transportability of 12 mouse phenotypes to matched human phenotypes (total cholesterol, triglycerides, albumin, bilirubin, alkaline phosphatase, lymphocyte counts, mean corpuscular volume, mean platelet volume, platelet counts, length/height, red blood cell width, bone mineral density) and some evidence for glucose tolerance-related mouse models and human diabetes mellitus. These findings indicate potential value of mouse knockout data for these phenotypes for understanding perturbation effects of orthologue genes on the corresponding phenotypes in humans.

On average mouse estimates were positively correlated with human phenotypes across the tested phenotypes but there was substantial phenotypic heterogeneity, and the degree of correlation was modest. We found limited evidence of transportability for more than half of the matched mouse phenotypes including fat mass, HDL cholesterol and BMI. There are several potential explanations. First, transportability estimates are context-specific and so our estimates should be interpreted in the context of the experimental conditions of the IMPC mice. The experimental conditions (early adulthood, standard diets, specific pathogen free environment) likely impact transportability to human phenotypes measured in late adulthood. Indeed, wild and laboratory mice have been previously shown to exhibit immunological differences with wild mice having higher immunoglobulin serum concentrations and different T cell profiles ^30^. Second, genuine interspecies differences between mice and humans relating to orthologous gene function and/or differences in genetic architecture. For example, mice and humans may have differential buffering or homeostatic mechanisms leading to intra-species heterogeneity in gene perturbation effects ^31^. Third, phenotyping differences between mice and humans. The constituents of composite phenotypes (e.g., lymphocyte counts or total cholesterol) may differ between humans and mice which will affect genetic architecture and transportability ^32^. Fourth, technical factors such as spurious outlier driven associations in mice, variant mis annotation, measurement errors (in mice or humans) or differences in covariates (some mouse estimates adjusted for weight). The IMPC is a high-throughput database which is not manually curated or measurement-repeated at an individual mouse level and so some datapoints may be erroneous. We used unweighted (but signed) mouse effect sizes in GPAT analyses because of difficulties standardising heterogeneous effect sizes across the IMPC database. As a result, the GPAT estimates are losing resolution by not modelling continuous magnitudes in the mouse data. Fifth, mouse models were a mixture of heterozygous and homozygous knockouts across different genes while the human estimates are based primarily on heterozygous pLoF. Non-linear effects of gene dosage could impact GPAT estimates. Sixth, we only included genes where the mouse gene had a high confidence ortholog in humans with a one-to-one mapping.

Directionally discordant effects have a large impact on GPAT estimates so we systematically explored their prevalence, finding that the majority of perturbation-phenotype associations observed in humans were directionally concordant with mouse models, but a substantial proportion were discordant. In particular we observed seven examples of perturbations with very strong associations in humans (P < 1x10^-5^) on mean platelet volume, alkaline phosphatase, eosinophil counts and mean corpuscular volume, where the mouse effect estimate was in the opposite direction. Manual review suggested that some of these discordant associations may relate to outliers in the mouse data warranting further investigation. In sensitivity analyses we found limited evidence of pLoF variant misannotation in humans for discordant pairs, and that base-pair sequence similarity was similar in discordant and concordant pairs. Further work is required to understand the impact of mouse-human gene orthology. In addition, these discordant results illustrate that while GPAT can indicate general relevance of a mouse phenotype to humans, it does not exclude species-specific differences in function of individual genes or pathways.

Improved understanding of directionally discordant effects is important to optimise the use of mouse knockout data in therapeutic development (e.g. modifying experimental conditions), avoid potentially misleading results and decision-making errors in drug development pipelines. For example, gene perturbation of *PCSK9* was strongly associated with lower HDL-cholesterol (HDL-C) in mice but, conversely, was positively associated with higher HDL-C in humans (P < 0.05) in the UK Biobank data. The human genetics evidence was consistent with the randomised trial of a *PCSK9* inhibitor which also observed elevated HDL-C in the treatment arm ^33^. Although there was a positive correlation for total cholesterol, effect estimates on HDL were not positively correlated between mice and humans indicative of cross-species aetiological heterogeneity, and these may reflect well known species differences in cholesterol metabolism in humans and mice, for example, absence of CETP in mice. A notable strength of GPAT over gene-set enrichment-based methods is that it models the direction of effect, not just the statistical significance of estimates. Directional information is highly desirable for informing possible interventions, such as a molecule lowering gene function, as otherwise the intervention may have an effect in the opposite-to-desired direction.

In previous work ^17^, we applied GPAT analyses to evaluate the transportability of *in-vitro* cellular models to human phenotypes and have now extended this approach to a large database of mouse knockout associations. Similar methods could be applied to any model organism, but mice are likely the best-studied, closely related model organism to humans. This could help provide more evidence about the transportability of these models for making inferences about human disease aetiology. The IMPC mouse genome database has partial coverage of human protein-coding genes with data on orthologues for just under half of human protein-coding genes. The database includes human genes with low pLoF frequency in UK Biobank, enabling it to provide valuable insights where human perturbation effects are highly imprecise. For some phenotypes, the limited GPAT evidence is likely due to the sparsity of IMPC mouse data, which currently does not include data on every phenotype in each knockout strain. For example, we observed very strong evidence of transportability for total cholesterol between mice and humans but with data on fewer genes (e.g. no *PCSK9* or *LDLR*) the matched LDL-cholesterol estimate was much more imprecise. A missing aspect of the IMPC mouse database is that it does not necessarily include well-characterised knockout relationships from the literature. For example, there is no data on the well-characterised *Mc4r* mouse model for obesity. Future work could complement this proof-of-concept analysis by systematically integrating literature evidence.

GPAT analyses use pLoF burden estimates as measures of effects of loss-of-function in humans. A previous study found evidence that > 25% of pLoF variants, such as those in the last exon, may not cause loss-of-function ^34^. GPAT estimates are reliant on accurate in-silico variant annotation and so may be biased towards the null. Similarly, IMPC knock-out strains have been generated by removing critical sequences for gene function over more than 15 years. Therefore, it is possible that some gene isoforms, not discovered at the time of null allele design, may be present in the mouse strains.

Our analysis was hypothesis-free, collapsing all genes according to the phenotype under assessment. This approach is necessarily scalable but leaves us less able to comment on component pathways and processes. There can be heterogeneity in the conservation of pathways regulating a phenotype of interest. For example, loss-of-function in components of the leptin-melanocortin system cause obesity in mice and humans via regulation of satiety – whereas differences in thermoneutral points between humans and mice mean peripheral mechanisms which regulate energy expenditure are less likely to replicate a-priori. The scaled approach cannot distinguish such nuance and is a limitation of note.

There is a substantial literature on mouse models being applied to understand complex human diseases via putative risk factors and endophenotypes such as insulin resistance (diabetes mellitus) alongside more direct approaches such as mouse tumour models ^35–40^. These cross-phenotype models rely on both the transportability of the mouse to matching human phenotype and causality between the human endophenotype and the disease. In an example of cross-phenotype GPAT analyses we found some evidence of transportability of results from dynamic glucose tolerance testing in mice to diabetes mellitus illustrating that GPAT analyses can validate mouse models for human diseases, in contrast to assessments of random glucose. These results have ethical as well as methodological implications, because they support the utilisation of intraperitoneal glucose tolerance testing, despite its relatively greater burden on mice compared to simpler spot tests such as fasting glucose. GPAT analyses could be particularly important for validating behavioural mouse models which are challenging to reliably link to human disease outcomes. pLoF burden tests of low prevalence diseases in humans are often underpowered with existing datasets so generating precise GPAT estimates is likely to require large disease-focused meta-analyses of rare variant gene analysis in humans ^5^. We also applied hypothesis-free cross-phenotype GPAT analyses to mouse total cholesterol finding some evidence of transportability to a number of human phenotypes beyond cholesterol such as triglycerides and adiposity phenotypes.

GPAT analyses provided strong evidence for transportability of a number of laboratory mouse phenotype models to corresponding human phenotypes with an overall positive, albeit modest, correlation observed between perturbation effects in mice and humans. However, we also observed interspecies heterogeneity in perturbation effects including instances of discordant directionality of effects between laboratory mice and humans. Further work is required to understand if such differences relate to genuine biological differences, lack of concordance between human lifestyle / experience and experimental mouse conditions, or technical errors relating to unharmonized data. We provide a framework for quantitatively validating experimental inbred mouse knockout models for understanding the aetiology of human phenotypes.

## Methods

### IMPC

The IMPC ^11^ makes available to the research community via a web portal (https://www.mousephenotype.org/) a large database of curated mouse gene knockout data including associations between gene knockout and hundreds of phenotypes. We downloaded data from v-24.0 of the database (https://ftp.ebi.ac.uk/pub/databases/impc/all-data-releases/release-24.0/) specifically downloading the file containing all results (statistical-results-ALL.csv.gz).

We manually matched mouse phenotypes (via MP: codes) with human phenotypes measured in UK Biobank by reviewing the mouse phenotypes with significant associations (P < 1x10^-5^) (https://www.informatics.jax.org/vocab/mp_ontology/) ^23^ and then determining if a matching phenotype was present in the UK Biobank data showcase (https://biobank.ndph.ox.ac.uk/showcase/). For example, ‘increased and decreased body length, MP:0001257/1258’ mouse phenotypes were paired with standing height in UK Biobank (data showcase ID: 50). The only mouse-human phenotype pairs which were not exactly matching were length (mouse) + height (human) and weight (mouse) + BMI (human). The length-height matching is a natural pairing while we selected weight as the most proximal measured mouse phenotype to BMI in humans. This manual matching resulted in 30 mouse-human phenotype pairs. Please see **Supplementary Table 1** for information on phenotype matching data between mice and humans.

We matched mouse genes with human ortholog genes (e.g. *Pcsk9* in mice and *PCSK9* in humans) by using high-confidence orthologues downloaded from Ensembl BioMart (GRCm39, downloaded November 20, 2023). High-confidence orthologues are called for pairs of mouse-human genes with 2_50% identical base-pair sequence (both human to mouse and mouse to human) and either a gene order conservation score 2_75 or genome alignment score 2_ 75. We also restricted to one-to-one mappings where a mouse gene is paired with only one human gene (and the human gene is paired with only one mouse gene).

Data required for GPAT analyses are the mouse gene being perturbed via knockdown, the phenotype, P-values and signed effect sizes. In some instances, there were duplicate rows for gene-phenotype pairs. For example, for total cholesterol prior to orthologue matching or P-value filtering, 414 genes had knockout data corresponding to 442 rows with 25 genes having two or more rows. For duplicate removal it was difficult at scale to determine which data point was the best to use. Therefore, we selected the row with the largest sample size (summing female and male mutants plus controls) for each gene.

There was substantial heterogeneity in effect estimates across different genes for the same phenotype. For example, absolute perturbation effect estimates (with P < 1x10^-5s^) on total cholesterol in mice ranged from 0.42 to 128 (mean 20.3, SD = 19.6).

A major source of heterogeneity is differential scaling of estimates with the web portal showing that some gene knockouts estimates were in terms of mg/dL and others mmol/L. To the best of our knowledge the dataset does not contain sufficient data to harmonise such unit discrepancies at scale. Other sources of heterogeneity include differences in experimental details such as the test centre, phenotype definitions and covariates. Notably some gene-phenotype estimates adjusted for weight while others did not. Such heterogeneity could be problematic for GPAT analyses which would be greatly affected by outliers. Therefore, we decided to use an unweighted approach of modelling the directionality but not the continuous effect sizes. i.e. across all data points, if a beta value is less than zero then set to -1 and if a beta value is greater than zero set to 1. This means that the GPAT analysis modelled only directionality in mice but both directionality and quantitative effect sizes in humans. Conceptually this would be similar to an unweighted (but signed) polygenic score.

### UK Biobank

UK Biobank is a population study of over 500,000 adult participants (40-69 years) who were recruited between 2006 and 2010 across the United Kingdom ^20^. Phenotype data on study participants includes anthropometric and biomarker measurements from baseline, questionnaire data and linked primary and secondary care electronic health record data. The majority of UK Biobank participants have whole genome sequence data ^21^ enabling whole-genome genome-wide association studies and aggregated rare variant tests collapsing rare and ultra rare variants by in-silico predicted function. The UK Biobank was approved by the North West Multi- centre Research Ethics Committee and all study participants gave informed consent.

For GPAT analyses we require estimates of effects of gene loss-of-function on human phenotypes to pair with mouse knockout effects. As discussed in previous work ^17^, we used pLoF burden tests because of the high-confidence that pLoF variants lower gene function and because burden tests provide signed effect sizes (while most other methods provide only P-values). We performed pLoF gene burden tests on 450K UK Biobank participants of non-Finnish European ancestry ^21^ using REGENIE (v3.4.1) ^41^. Burden tests were performed using a mask of high-confidence LoF variants (based on variant-effect predictions: VEP) with MAF < 0.01. The default REGENIE burden model was used which assigns individuals values of 0 (no heterozygous or homozygous pLoF variants), 1 (one or more heterozygous pLoF variants) or 2 (one or more homozygous pLoF variants). pLoF burden data were available on up to 18,063 protein-coding genes which had 1 or more pLoF carrier in the whole cohort.

For quantitative phenotypes, we included 31 phenotypes (30 matched with mouse phenotypes and Hba1c) from the blood and urine biochemistry biomarker phenotypes (UK Biobank Showcase Category 17518), anthropometric phenotypes from the baseline assessment data (UK Biobank Showcase Category 100010) and plasma cell count phenotypes (UK Biobank Showcase Category 100081) (**Supplementary Table 1**). Diabetes mellitus was defined based on a combination of electronic health record data sources from CALIBER atlas phenotypes ^42^. An additional quantitative phenotype TG:HDL ratio was computed as the ratio of triglycerides and HDL-cholesterol. Quantitative phenotypes were rank-based inverse-normal transformed prior to analysis.

For proteomics, 2923 unique proteins were measured in plasma from 54,219 UKB participants using the antibody-based Olink Explore 3072 PEA. These proteins were then matched to the corresponding gene encoding the protein ^43^. In analyses, we started with pLoF burden data on 2,121 proteins which had 1 or more pLoF carriers in the study sample and then subset this down to a small set of 109 discordant genes with discordant associations between mice and humans.

### GPAT analyses

To combine mouse and human perturbation data for GPAT analyses we performed the following QC and harmonisation steps:

1. Remove duplicates in mouse knockout database for gene-phenotype pairs by taking the row with the largest sample size. This step is performed prior to the second step in order to prioritise the most reliable estimates.
2. Restrict to mouse gene-phenotype pairs with P < 1x10^-5^ and set mouse perturbation effect sizes to 1 or -1 depending on directionality of effect.
3. Merge the mouse dataset with the UK Biobank pLoF burden data of the matched phenotype (e.g. mouse length and human height) by merging on orthology-matched gene symbol. We restricted to orthologous pairs of high confidence with one-to-one mappings. This step would also remove a small number of genes with no pLoF carriers in UK Biobank.

We used linear regression models in R to assess the association between pLoF cumulative frequency and available matched mouse knockout data across the full set of mouse genes represented in the IMPC database (which are matched with orthologous human genes) and subset of mouse genes with perturbation evidence for one or more mouse phenotypes (3483 genes, P < 1x10^-5^).

As described above, the GPAT framework is a gene-level implementation of Mendelian randomization. Wald ratios are calculated for each gene using the perturbation effect sizes on the phenotype (*β_j,/_, β_j,k_)* and corresponding standard errors.

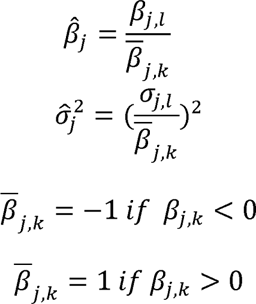

We calculated Wald ratios using the equations above and then used the R package *metagen* to perform fixed-effects and random-effects inverse-variance weighted meta-analyses and generate funnel plots. We reported the fixed-effects estimates as the primary analysis with cross-reference of the random-effects estimates.

We performed same-phenotype GPAT analyses for the matched mouse-human phenotypes (30 phenotypes in total). For example, total cholesterol in mice and total cholesterol in humans. With a strong prior for transportability due to the same phenotype being considered in both mice and humans we use a liberal P-value threshold of 0.05 for the reporting of results with no correction for multiple testing. For the same-phenotype GPAT analyses we meta-analysed the individual GPAT estimates for the 30 phenotypes to generate an estimate of the average transportability across all tested phenotypes. With the caveat that some of the tested phenotypes are correlated (e.g. LDL-cholesterol and total cholesterol) this approach enables an estimate of the average correlation across the tested phenotypes. We performed cross-phenotype GPAT analyses for total-cholesterol in mice and the remaining 29 phenotypes in humans.

For the diabetes mellitus analysis, we defined the five mouse gene sets as follows:

1. Glucose tolerance based on the area under a glucose response curve in an Intraperitoneal glucose tolerance test: IPGTT corresponding to improved glucose tolerance, impaired glucose tolerance and abnormal glucose homeostasis (MP: 0005292, MP:0005293, MP:0002078). Note that higher values of IPGTT correspond to impaired glucose tolerance linked to higher risk of diabetes.
2. Fasting glucose as measured in an IPGTT after eating corresponding to decreased and increased fasting circulating glucose levels (MP:0013278; MP:0013279).
3. Free-fed glucose as measured in an IPGTT corresponding to decreased and increased circulating glucose levels (MP:0005559, MP:0005560).
4. Glucose all is a union of mouse knockouts associated with glucose tolerance, fasting glucose and free-fed glucose (MP: 0005292, MP:0005293, MP:0002078, MP:0013278, MP:0013279, MP:0005559, MP:0005560). If a gene is in both of the sets then the most precise effect size is prioritised for the directionality.
5. Insulin as measured in the blood corresponding to increased or decreased circulating levels (MP:0002079; MP:0002727).

We then performed cross-trait GPAT analyses using these five gene-sets with diabetes mellitus, Hba1c and TG:HDL ratio in humans.

Outlier genes were visually inspected from GPAT funnel plots. In alkaline phosphatase analyses, we removed *GPLD1* because of the directional discordance and extremely strong association in humans. Indeed, there was substantial heterogeneity between the fixed-effects and random-effects GPAT estimates before excluding *GPLD1*. In specific instances we also manually reviewed scatter plots of individual-level data from the mouse database web application (https://www.mousephenotype.org/) to better understand some gene-phenotype observations in mice. For example, the association between *RASAL3* knockdown and higher eosinophil differential count seemed to be observed only in two female mutants.

To investigate discordance between mouse and human perturbation estimates in same-phenotype associations, we defined two sets of gene-phenotype pairs with evidence in mice (P < 1x10^-5^) and in humans (pLoF burden P < 0.05 or pLoF burden P < 1x10^-5^). These pairs were then defined as concordant or discordant depending on sign similarity of effect sizes. Merging these data with the proteomics data we calculated the median and quartiles using R functions. We used linear regression in R to evaluate the association between base-pair sequence similarity and concordance.

### Mouse sign concordance analyses

We wanted to evaluate if genes where mouse knockouts are strongly associated (P < 1x10^-5^) with total cholesterol are also associated with other murine phenotypes with consistent directionality. However, the challenges with running a GPAT-style analysis are two-fold. First, as discussed, the mouse effect size estimates are heterogeneous and difficult to standardise. Second, there is sparse coverage of gene-phenotype data within the database with only a subset of genes having data for each phenotype.

Therefore, we focused on concordance in sign (without modelling effect sizes) to use all available data points. To reduce sensitivity to low counts we used a binomial sign test. This involved joining the 131 genes for total cholesterol (P < 1x10^-5^) with available data for each of the 29 murine phenotypes. The number of genes is higher than for the mouse-human analyses because orthologue matching was not required. The binomial sign test was performed on the numbers of concordant and discordant genes based on direction of effects. We used a liberal P-value threshold of P < 0.05 to identify murine phenotypes where there is some evidence of more directional consistency than would be expected by chance.

## Acknowledgements

GDS works within the MRC Integrative Epidemiology Unit at the University of Bristol, which is supported by the Medical Research Council (MC_UU_00032/1). SW works within the Mary Lyon Centre at MRC Harwell which is supported by the Medical Research Council (MC_UP_2201/1) and the Biological Research Facility at the Francis Crick Institute.

## Author contributions

LJH conceived the project, performed all analyses and drafted the manuscript. LJH, GDS, SW, NMD and SML reviewed and critically contributed to the manuscript.

## Supplementary Figures

**Supplementary Figure 1.**
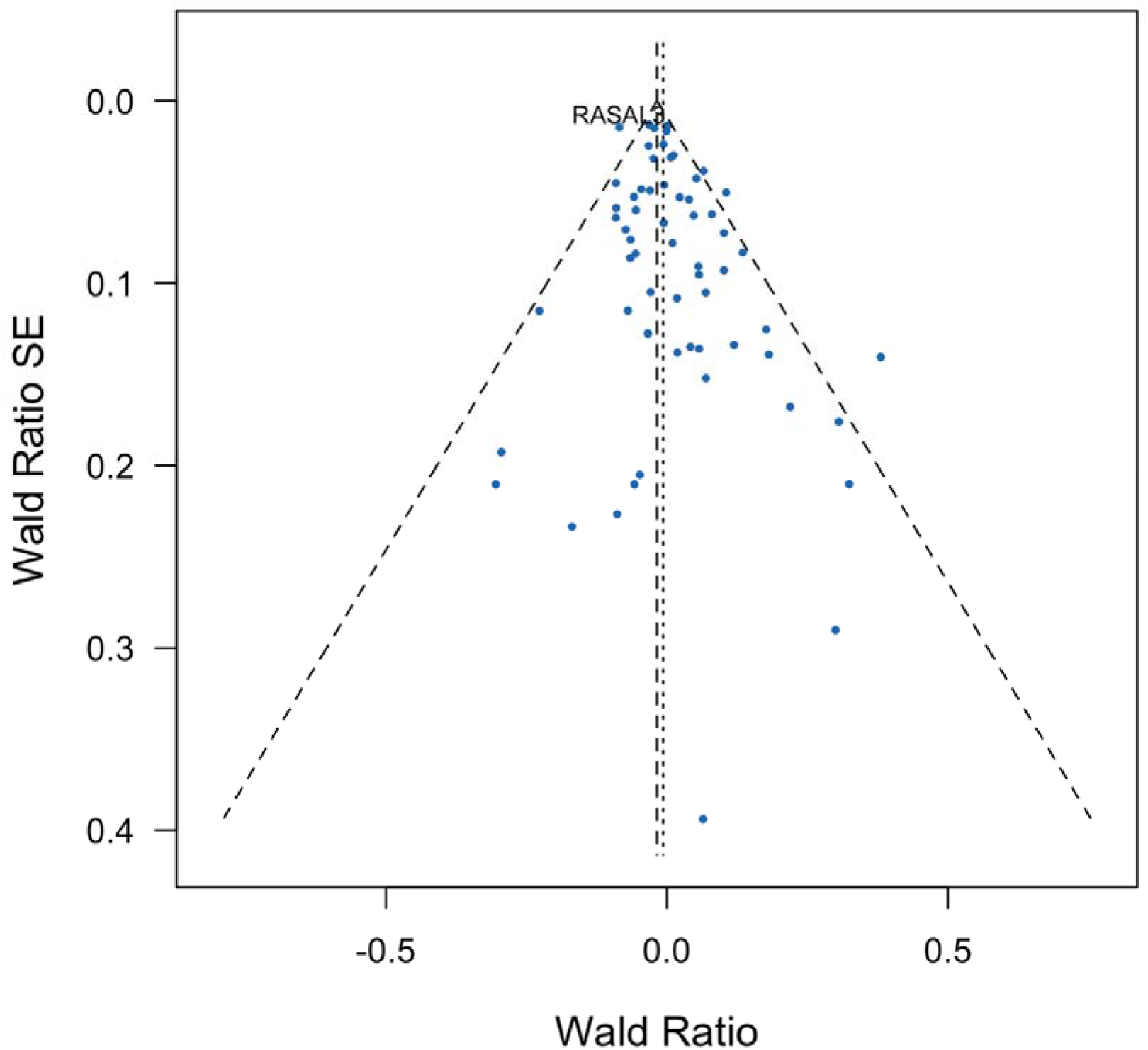
Gene perturbations strongly associated with eosinophil counts in mice and their association with eosinophils in humans Supplementary Figure 1 presents Wald ratios and standard errors for individual genes strongly associated with eosinophil counts in mice. Wald ratios are the gene-phenotype effect estimate in humans divided by the gene-phenotype effect estimate in mice. We used directional, unweighted mouse estimates and so the Wald ratios are the effect estimates from humans with positive (consistent directionality between mice and humans) or negative (inconsistent directionality) values. The dotted vertical lines indicate the overall fixed and random effects GPAT estimates across all genes. Some notable genes strongly associated with eosinophil counts in humans are manually annotated. All genes with pLoF burden P < 0.001 in humans are annotated with the human gene symbol. These estimates model only direction of effect in mice and both direction of effect and magnitude of effect in humans.

**Supplementary Figure 2.**
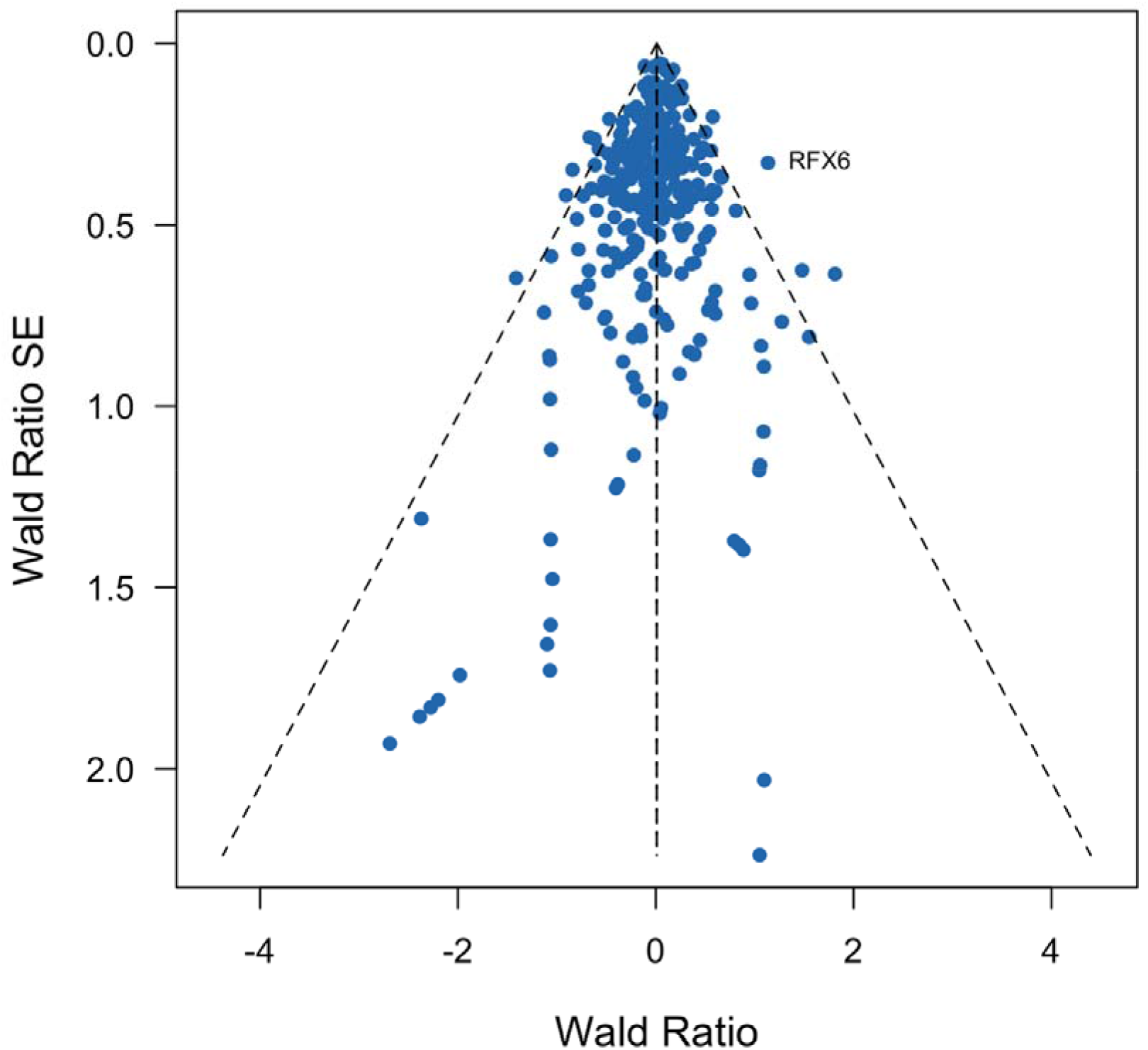
Gene perturbations strongly associated with glucose all in mice and their association with diabetes mellitus in humans Supplementary Figure 2 presents Wald ratios and standard errors for individual genes strongly associated with glucose all (fasting and/or tolerance) in mice with respect to diabetes mellitus in humans. Wald ratios are the gene-phenotype effect estimate in humans divided by the gene-phenotype effect estimate in mice. We used directional, unweighted mouse estimates and so the Wald ratios are the effect estimates from humans with positive (consistent directionality between mice and humans) or negative (inconsistent directionality) values. The dotted vertical lines indicate the overall fixed and random effects GPAT estimates across all genes. All genes with pLoF burden P < 0.001 in humans are annotated with the human gene symbol. These estimates model only direction of effect in mice and both direction of effect and magnitude of effect in humans.

**Supplementary Figure 3.**
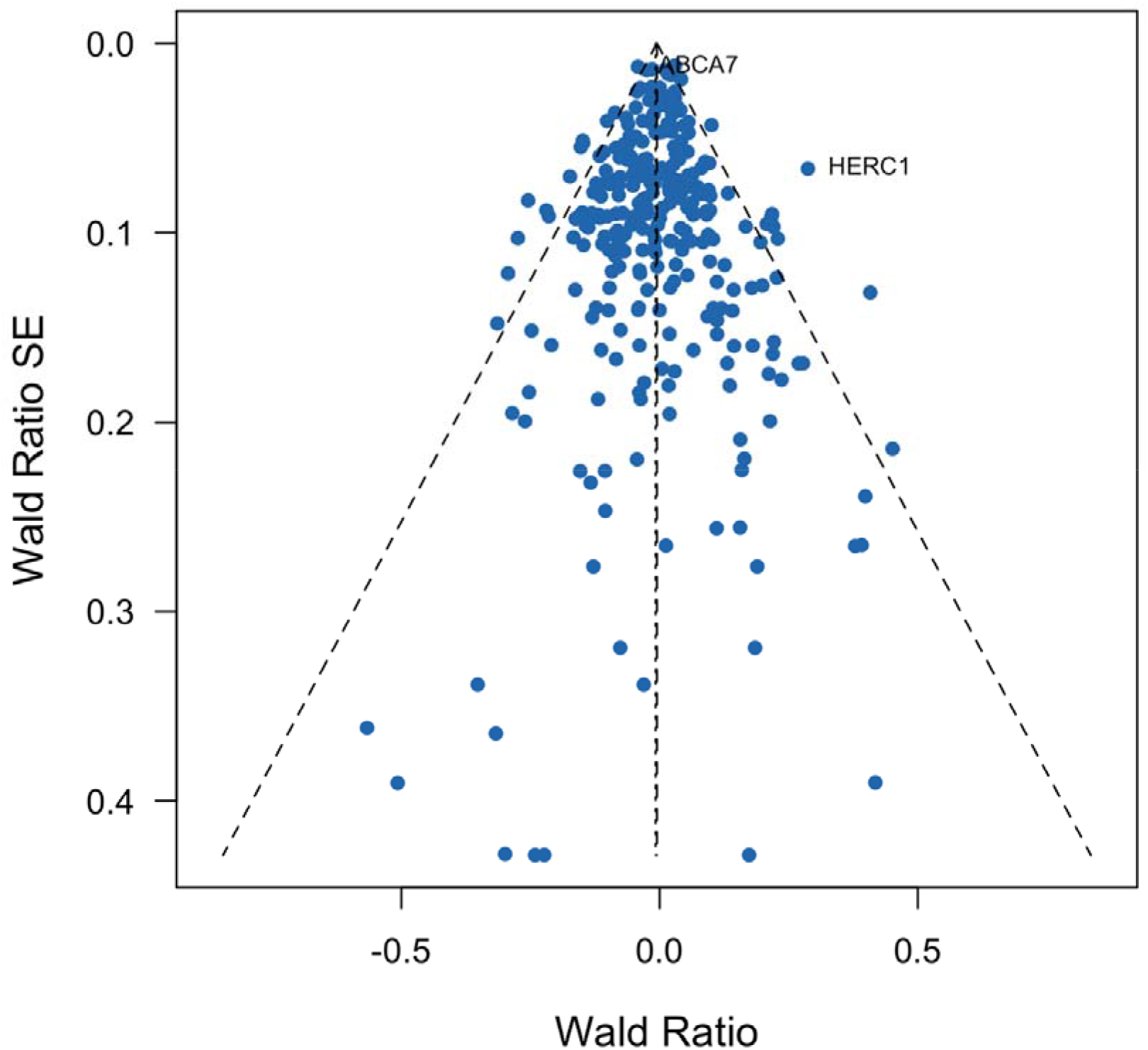
Gene perturbations strongly associated with glucose all in mice and their association with Hba1c in humans Supplementary Figure 3 presents Wald ratios and standard errors for individual genes strongly associated with glucose all (fasting and/or tolerance) in mice with respect to Hba1c in humans. Wald ratios are the gene-phenotype effect estimate in humans divided by the gene-phenotype effect estimate in mice. We used directional, unweighted mouse estimates and so the Wald ratios are the effect estimates from humans with positive (consistent directionality between mice and humans) or negative (inconsistent directionality) values. The dotted vertical lines indicate the overall fixed and random effects GPAT estimates across all genes. All genes with pLoF burden P < 0.001 in humans are annotated with the human gene symbol. These estimates model only direction of effect in mice and both direction of effect and magnitude of effect in humans.

**Supplementary Figure 4.**
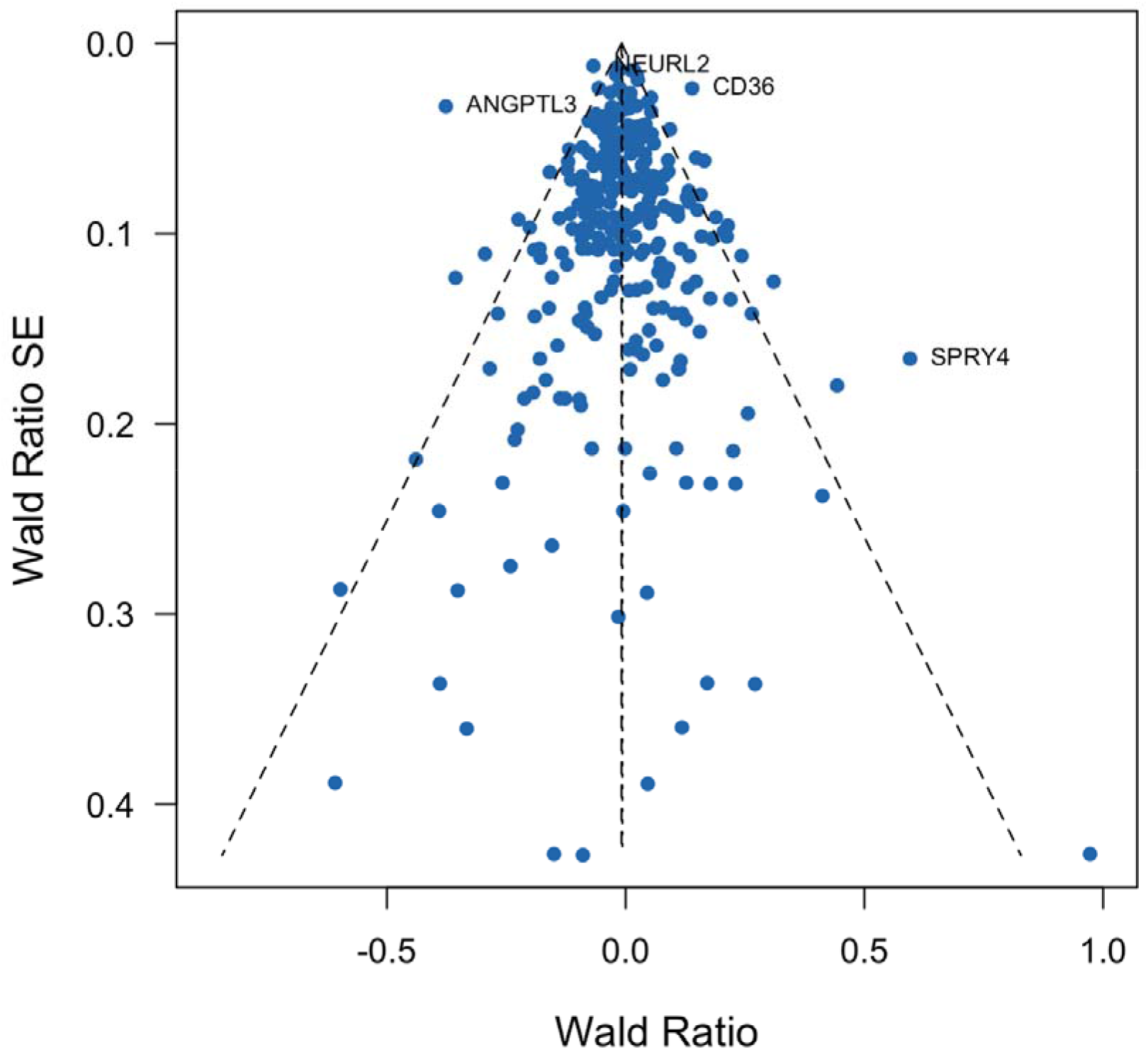
Gene perturbations strongly associated with glucose all in mice and their association with TG:HDL in humans Supplementary Figure 4 presents Wald ratios and standard errors for individual genes strongly associated with glucose all (fasting and/or tolerance) in mice with respect to TG:HDL in humans. Wald ratios are the gene-phenotype effect estimate in humans divided by the gene-phenotype effect estimate in mice. We used directional, unweighted mouse estimates and so the Wald ratios are the effect estimates from humans with positive (consistent directionality between mice and humans) or negative (inconsistent directionality) values. The dotted vertical lines indicate the overall fixed and random effects GPAT estimates across all genes. All genes with pLoF burden P < 0.001 in humans are annotated with the human gene symbol. These estimates model only direction of effect in mice and both direction of effect and magnitude of effect in humans.

**Supplementary Figure 5.**
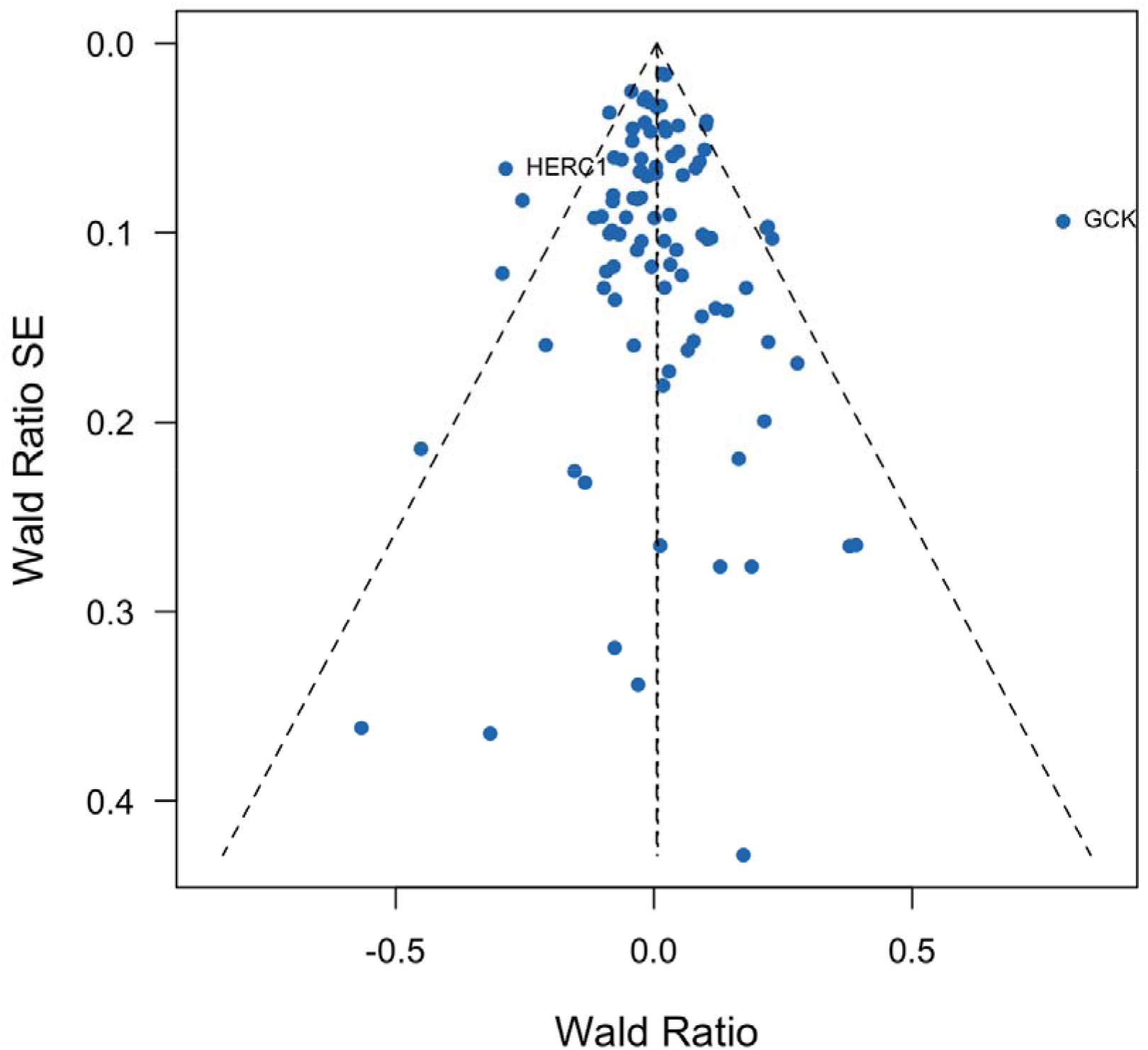
Gene perturbations strongly associated with glucose tolerance in mice and their association with Hba1c in humans Supplementary Figure 5 presents Wald ratios and standard errors for individual genes strongly associated with glucose tolerance in mice with respect to Hba1c in humans. Wald ratios are the gene-phenotype effect estimate in humans divided by the gene-phenotype effect estimate in mice. We used directional, unweighted mouse estimates and so the Wald ratios are the effect estimates from humans with positive (consistent directionality between mice and humans) or negative (inconsistent directionality) values. The dotted vertical lines indicate the overall fixed and random effects GPAT estimates across all genes. All genes with pLoF burden P < 0.001 in humans are annotated with the human gene symbol. These estimates model only direction of effect in mice and both direction of effect and magnitude of effect in humans.

**Supplementary Figure 6.**
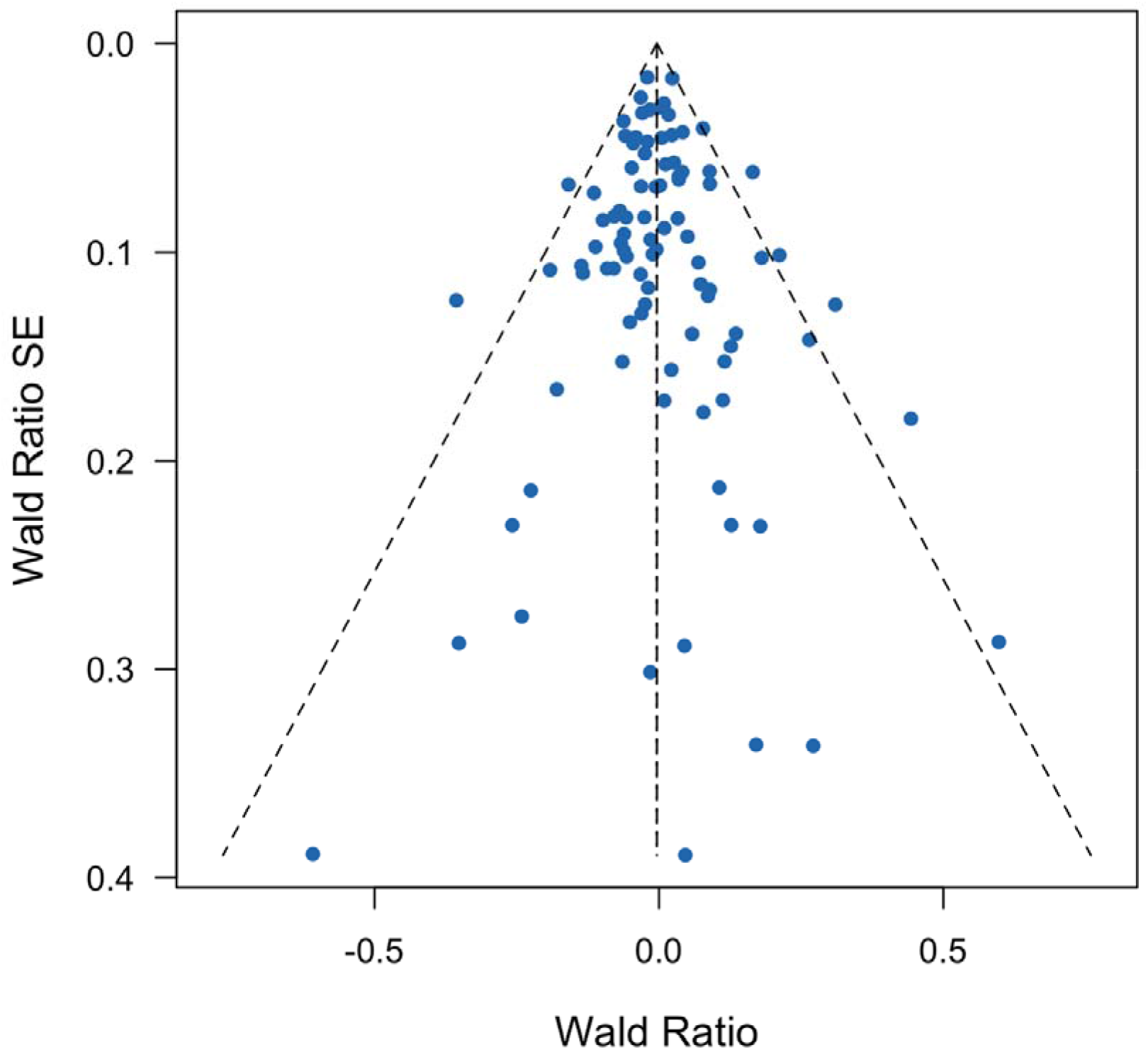
Gene perturbations strongly associated with glucose tolerance in mice and their association with TG:HDL in humans Supplementary Figure 6 presents Wald ratios and standard errors for individual genes strongly associated with glucose tolerance in mice with respect to TG:HDL in humans. Wald ratios are the gene-phenotype effect estimate in humans divided by the gene-phenotype effect estimate in mice. We used directional, unweighted mouse estimates and so the Wald ratios are the effect estimates from humans with positive (consistent directionality between mice and humans) or negative (inconsistent directionality) values. The dotted vertical lines indicate the overall fixed and random effects GPAT estimates across all genes. All genes with pLoF burden P < 0.001 in humans are annotated with the human gene symbol. These estimates model only direction of effect in mice and both direction of effect and magnitude of effect in humans. These estimates model only direction of effect in mice and both direction of effect and magnitude of effect in humans.

**Supplementary Figure 7.**
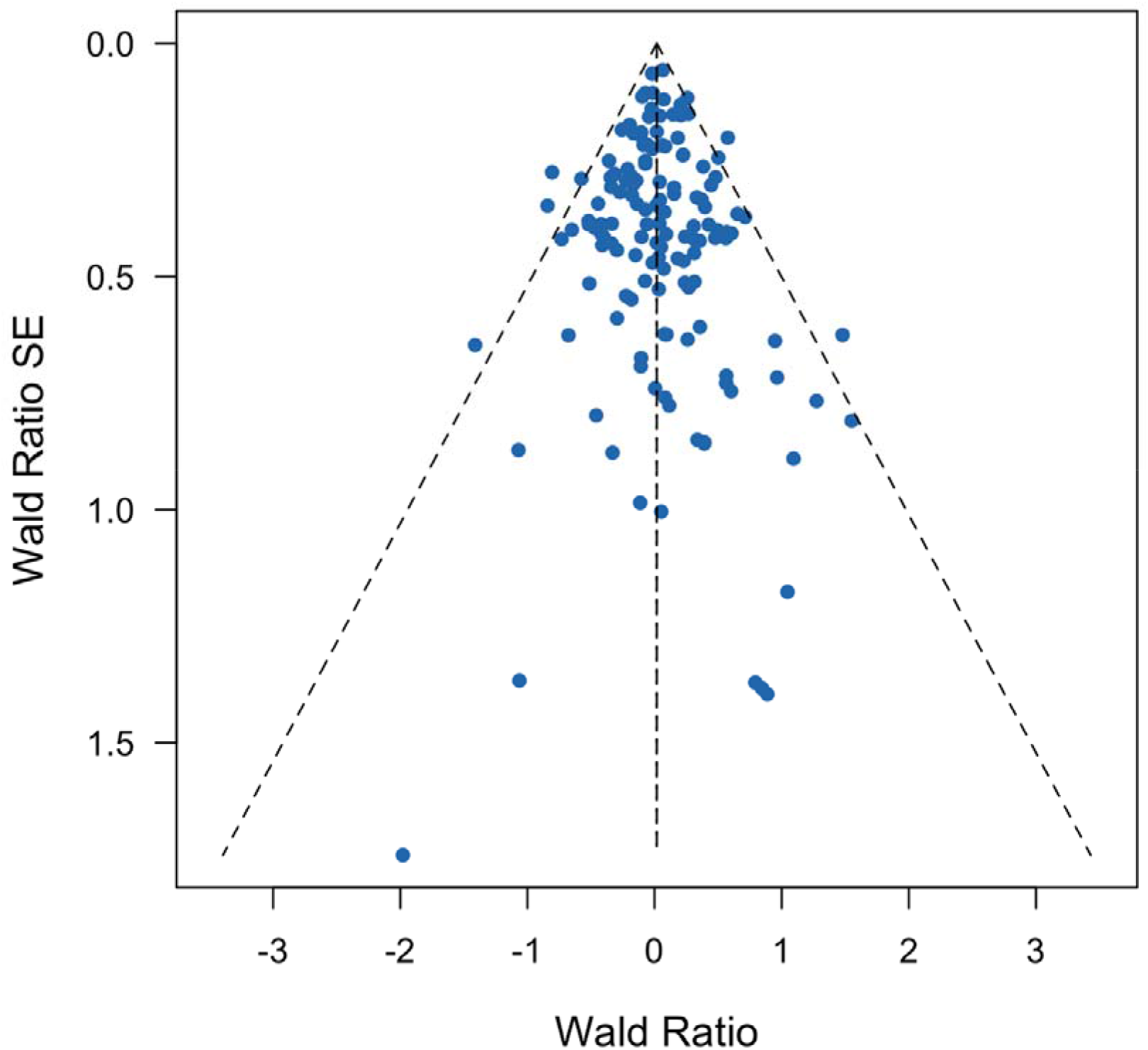
Gene perturbations strongly associated with fasting glucose in mice and their association with diabetes mellitus in humans Supplementary Figure 7 presents Wald ratios and standard errors for individual genes strongly associated with fasting glucose in mice with respect to diabetes mellitus in humans. Wald ratios are the gene-phenotype effect estimate in humans divided by the gene-phenotype effect estimate in mice. We used directional, unweighted mouse estimates and so the Wald ratios are the effect estimates from humans with positive (consistent directionality between mice and humans) or negative (inconsistent directionality) values. The dotted vertical lines indicate the overall fixed and random effects GPAT estimates across all genes. All genes with pLoF burden P < 0.001 in humans are annotated with the human gene symbol. These estimates model only direction of effect in mice and both direction of effect and magnitude of effect in humans.

**Supplementary Figure 8.**
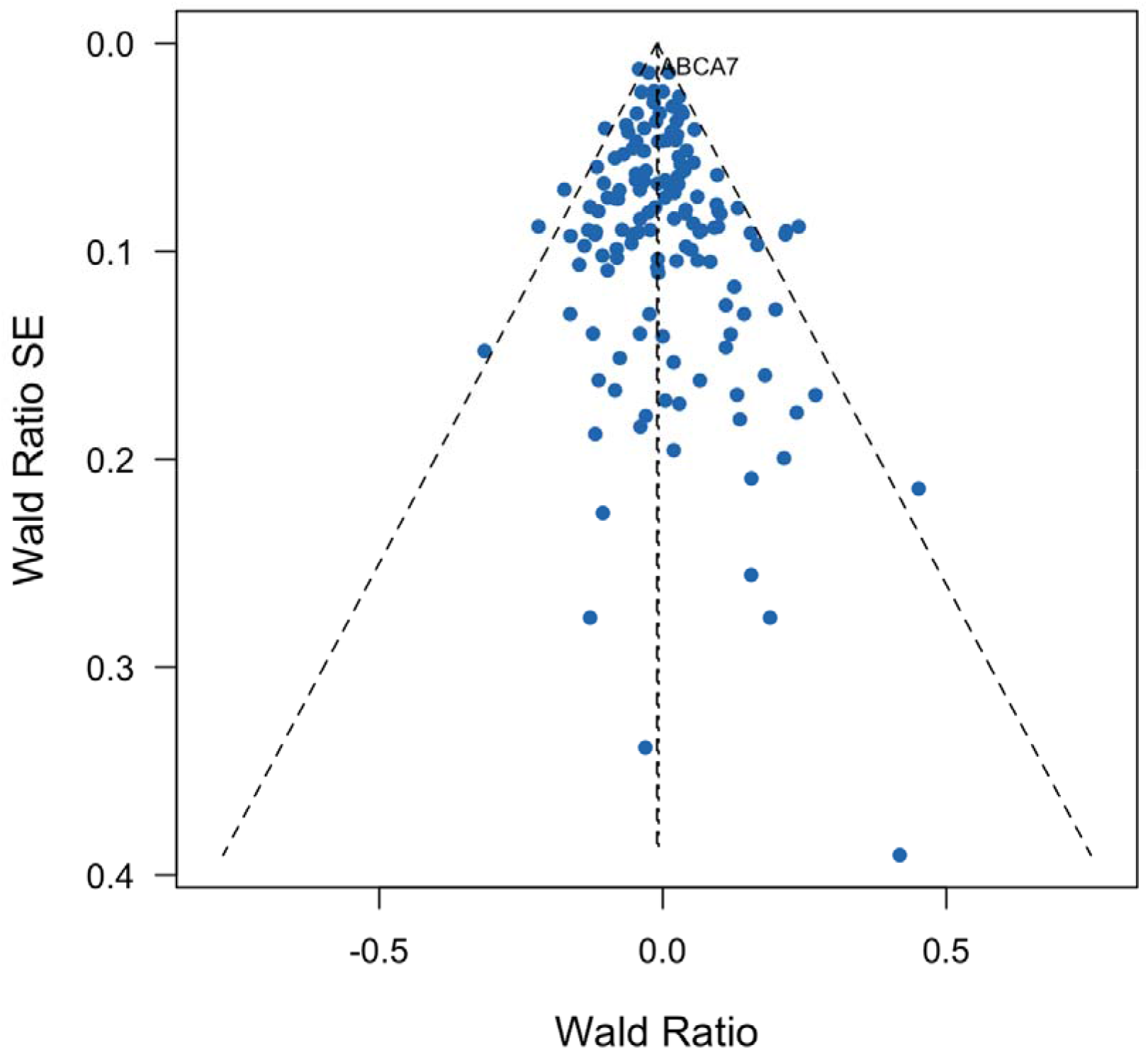
Gene perturbations strongly associated with fasting glucose in mice and their association with Hba1c in humans Supplementary Figure 8 presents Wald ratios and standard errors for individual genes strongly associated with fasting glucose in mice with respect to Hba1c in humans. Wald ratios are the gene-phenotype effect estimate in humans divided by the gene-phenotype effect estimate in mice. We used directional, unweighted mouse estimates and so the Wald ratios are the effect estimates from humans with positive (consistent directionality between mice and humans) or negative (inconsistent directionality) values. The dotted vertical lines indicate the overall fixed and random effects GPAT estimates across all genes. All genes with pLoF burden P < 0.001 in humans are annotated with the human gene symbol.

**Supplementary Figure 9.**
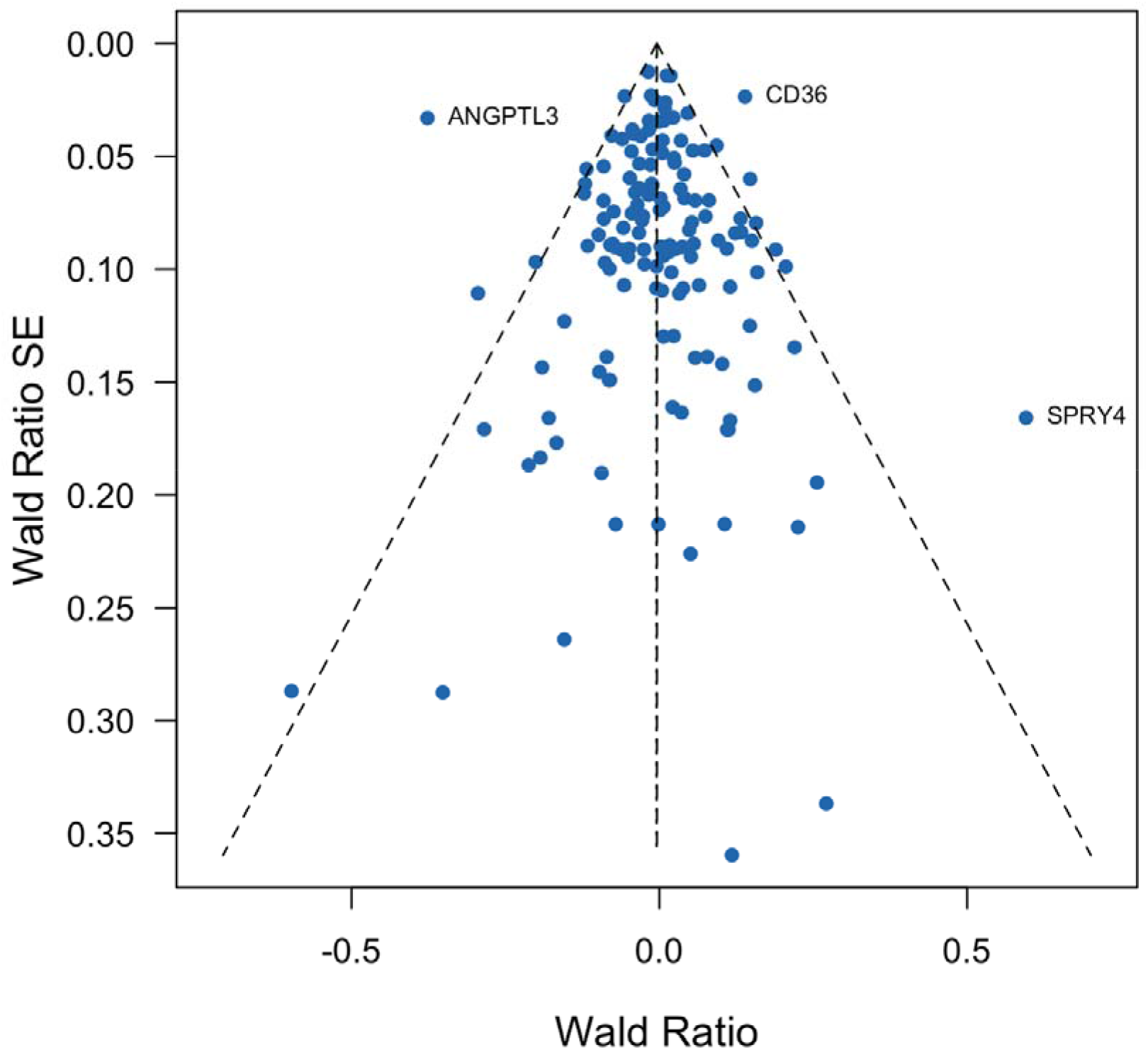
Gene perturbations strongly associated with fasting glucose in mice and their association with TG:HDL in humans Supplementary Figure 9 presents Wald ratios and standard errors for individual genes strongly associated with fasting glucose in mice with respect to TG:HDL in humans. Wald ratios are the gene-phenotype effect estimate in humans divided by the gene-phenotype effect estimate in mice. We used directional, unweighted mouse estimates and so the Wald ratios are the effect estimates from humans with positive (consistent directionality between mice and humans) or negative (inconsistent directionality) values. The dotted vertical lines indicate the overall fixed and random effects GPAT estimates across all genes. All genes with pLoF burden P < 0.001 in humans are annotated with the human gene symbol.

**Supplementary Figure 10.**
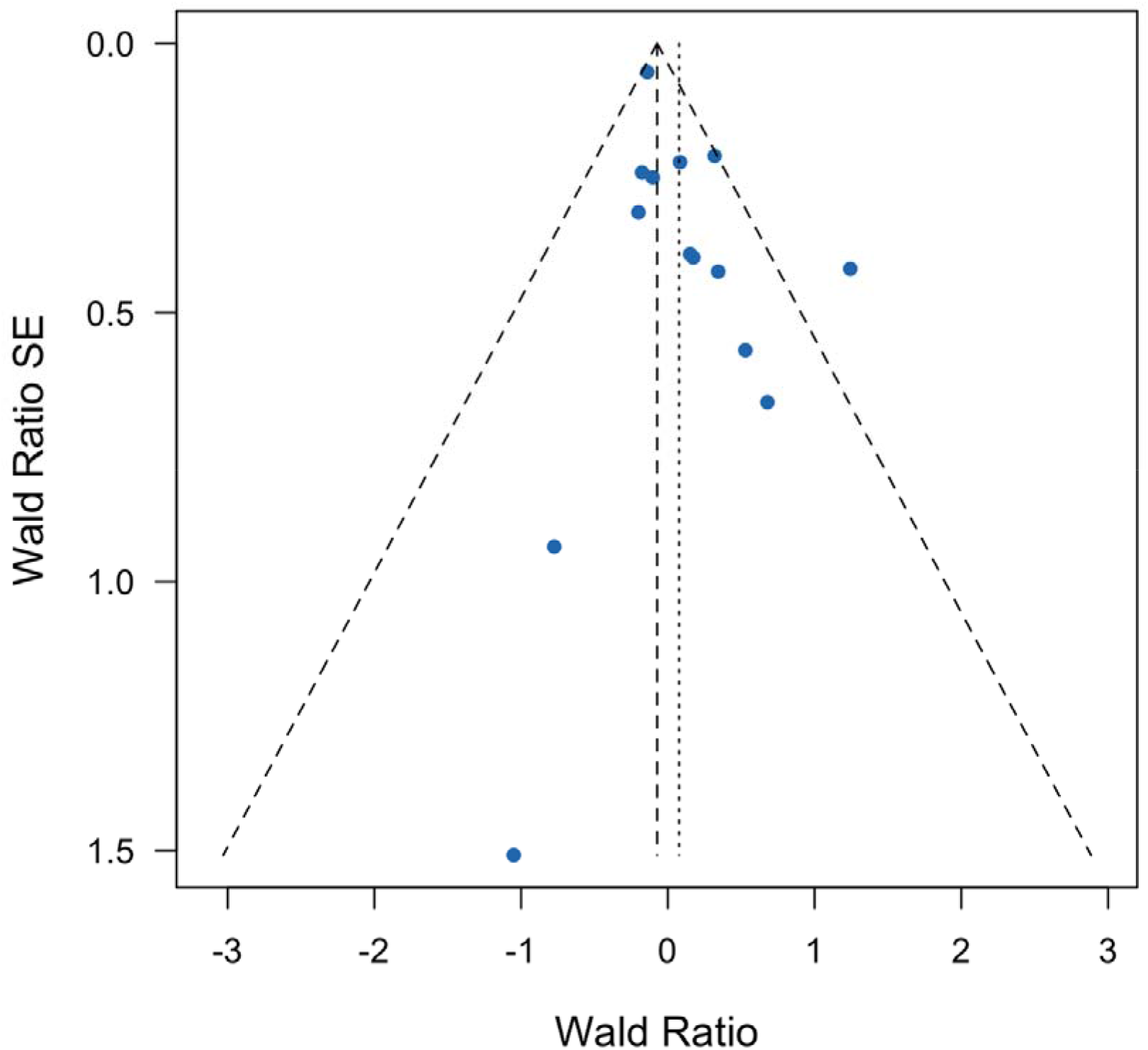
Gene perturbations strongly associated with circulating insulin in mice and their association with diabetes mellitus in humans Supplementary Figure 10 presents Wald ratios and standard errors for individual genes strongly associated with circulating insulin in mice with respect to diabetes mellitus in humans. Wald ratios are the gene-phenotype effect estimate in humans divided by the gene-phenotype effect estimate in mice. We used directional, unweighted mouse estimates and so the Wald ratios are the effect estimates from humans with positive (consistent directionality between mice and humans) or negative (inconsistent directionality) values. The dotted vertical lines indicate the overall fixed and random effects GPAT estimates across all genes. All genes with pLoF burden P < 0.001 in humans are annotated with the human gene symbol.

**Supplementary Figure 11.**
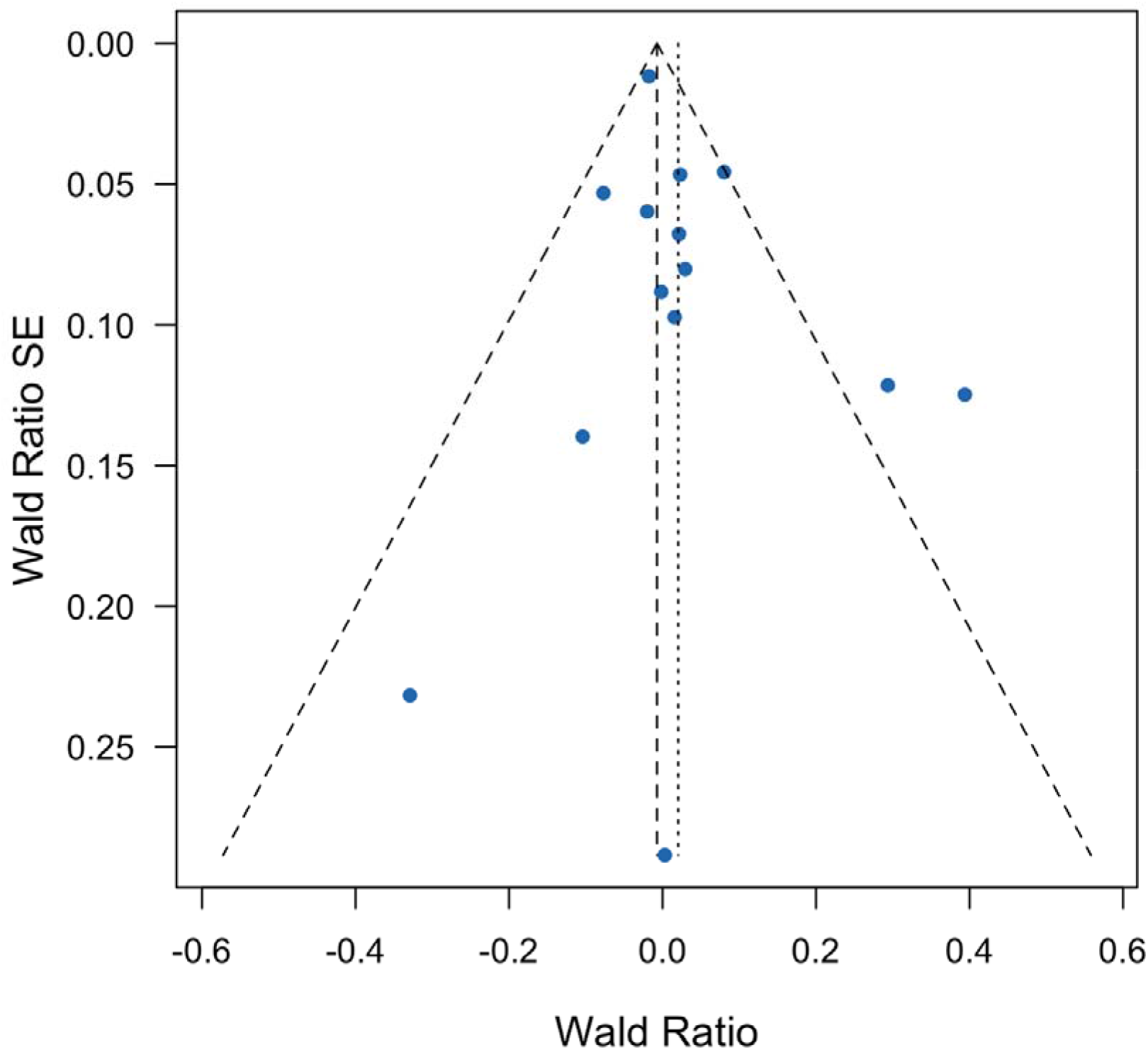
Gene perturbations strongly associated with circulating insulin in mice and their association with Hba1c in humans Supplementary Figure 11 presents Wald ratios and standard errors for individual genes strongly associated with circulating insulin in mice with respect to Hba1c in humans. Wald ratios are the gene-phenotype effect estimate in humans divided by the gene-phenotype effect estimate in mice. We used directional, unweighted mouse estimates and so the Wald ratios are the effect estimates from humans with positive (consistent directionality between mice and humans) or negative (inconsistent directionality) values. The dotted vertical lines indicate the overall fixed and random effects GPAT estimates across all genes. All genes with pLoF burden P < 0.001 in humans are annotated with the human gene symbol.

**Supplementary Figure 12.**
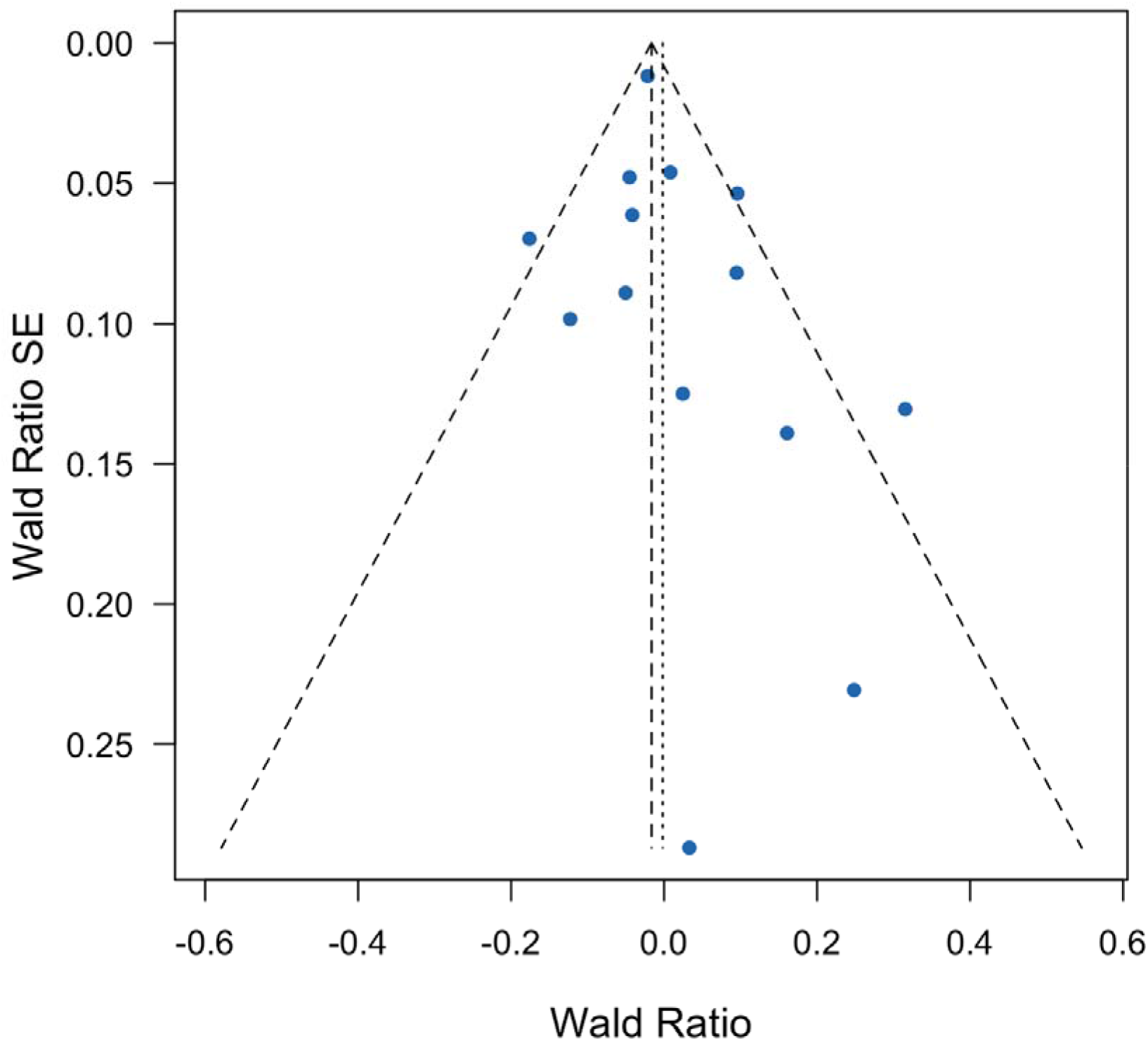
Gene perturbations strongly associated with circulating insulin in mice and their association with TG:HDL in humans Supplementary Figure 12 presents Wald ratios and standard errors for individual genes strongly associated with circulating insulin in mice with respect to TG:HDL in humans. Wald ratios are the gene-phenotype effect estimate in humans divided by the gene-phenotype effect estimate in mice. We used directional, unweighted mouse estimates and so the Wald ratios are the effect estimates from humans with positive (consistent directionality between mice and humans) or negative (inconsistent directionality) values. The dotted vertical lines indicate the overall fixed and random effects GPAT estimates across all genes. All genes with pLoF burden P < 0.001 in humans are annotated with the human gene symbol.

**Supplementary Figure 13.**
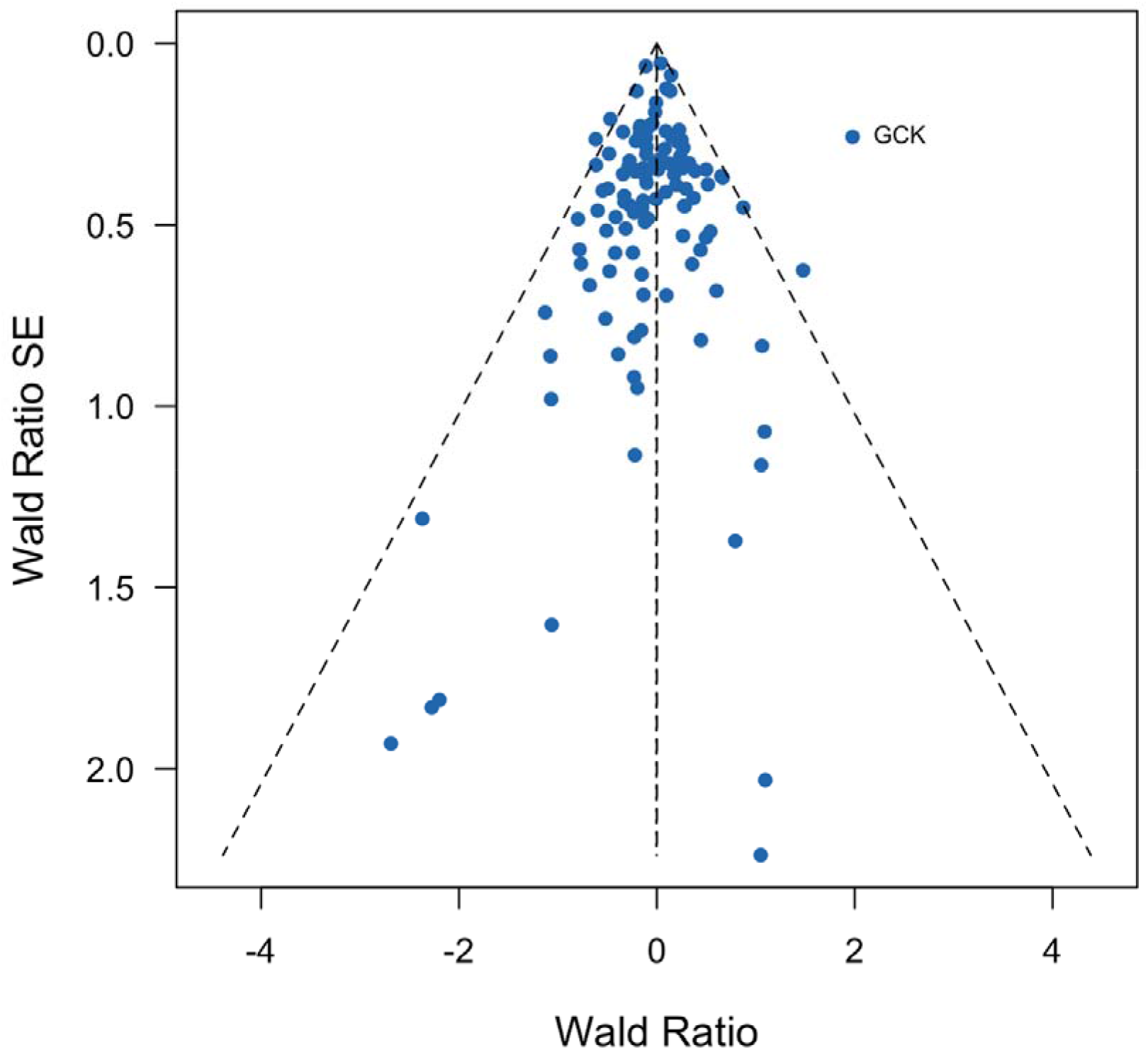
Gene perturbations strongly associated with free-fed glucose in mice and their association with diabetes mellitus in humans Supplementary Figure 13 presents Wald ratios and standard errors for individual genes strongly associated with free-fed glucose in mice with respect to diabetes mellitus in humans. Wald ratios are the gene-phenotype effect estimate in humans divided by the gene-phenotype effect estimate in mice. We used directional, unweighted mouse estimates and so the Wald ratios are the effect estimates from humans with positive (consistent directionality between mice and humans) or negative (inconsistent directionality) values. The dotted vertical lines indicate the overall fixed and random effects GPAT estimates across all genes. All genes with pLoF burden P < 0.001 in humans are annotated with the human gene symbol.

**Supplementary Figure 14.**
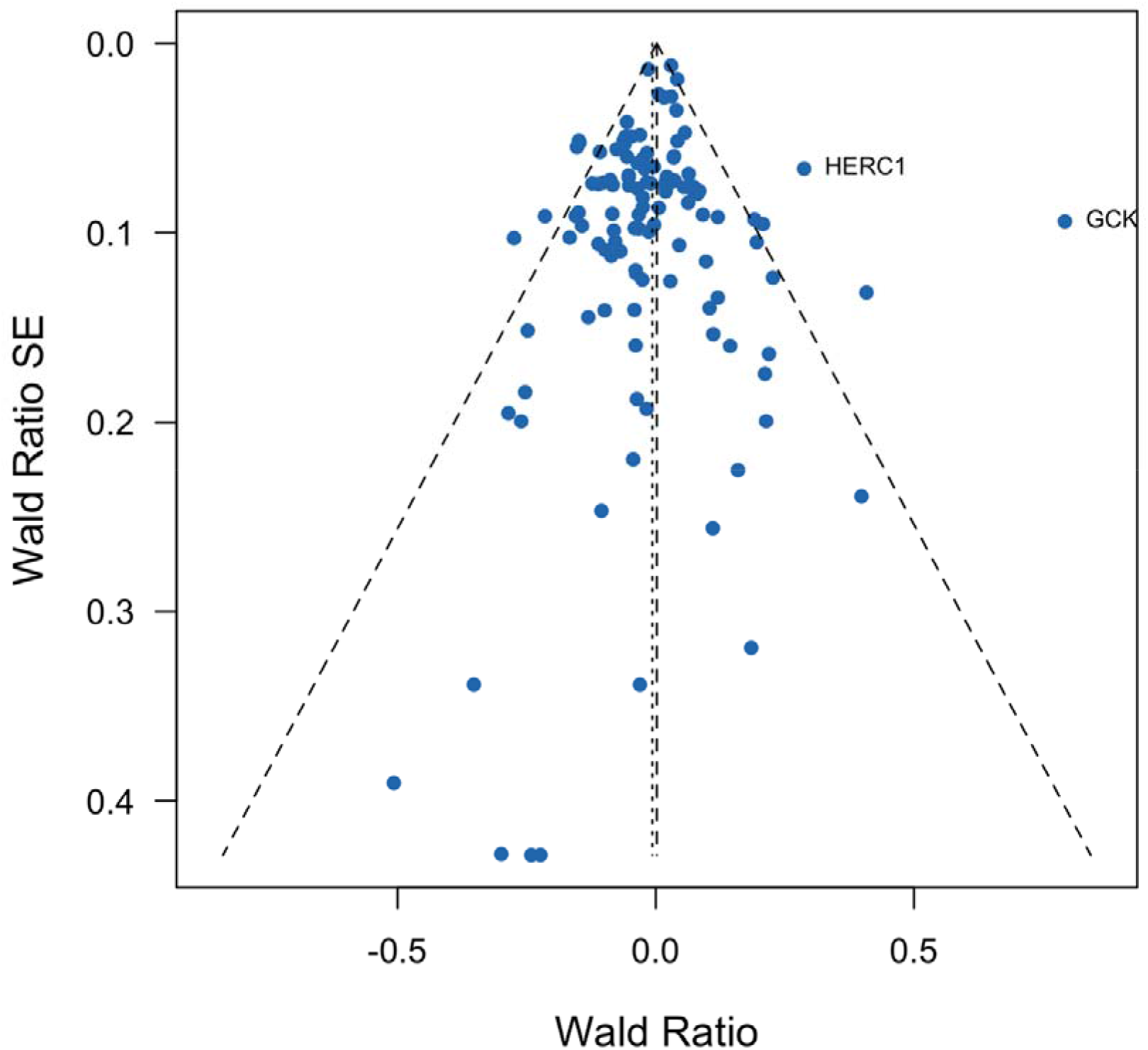
Gene perturbations strongly associated with free-fed glucose in mice and their association with Hba1c in humans Supplementary Figure 14 presents Wald ratios and standard errors for individual genes strongly associated with free-fed glucose in mice with respect to Hba1c in humans. Wald ratios are the gene-phenotype effect estimate in humans divided by the gene-phenotype effect estimate in mice. We used directional, unweighted mouse estimates and so the Wald ratios are the effect estimates from humans with positive (consistent directionality between mice and humans) or negative (inconsistent directionality) values. The dotted vertical lines indicate the overall fixed and random effects GPAT estimates across all genes. All genes with pLoF burden P < 0.001 in humans are annotated with the human gene symbol.

**Supplementary Figure 15.**
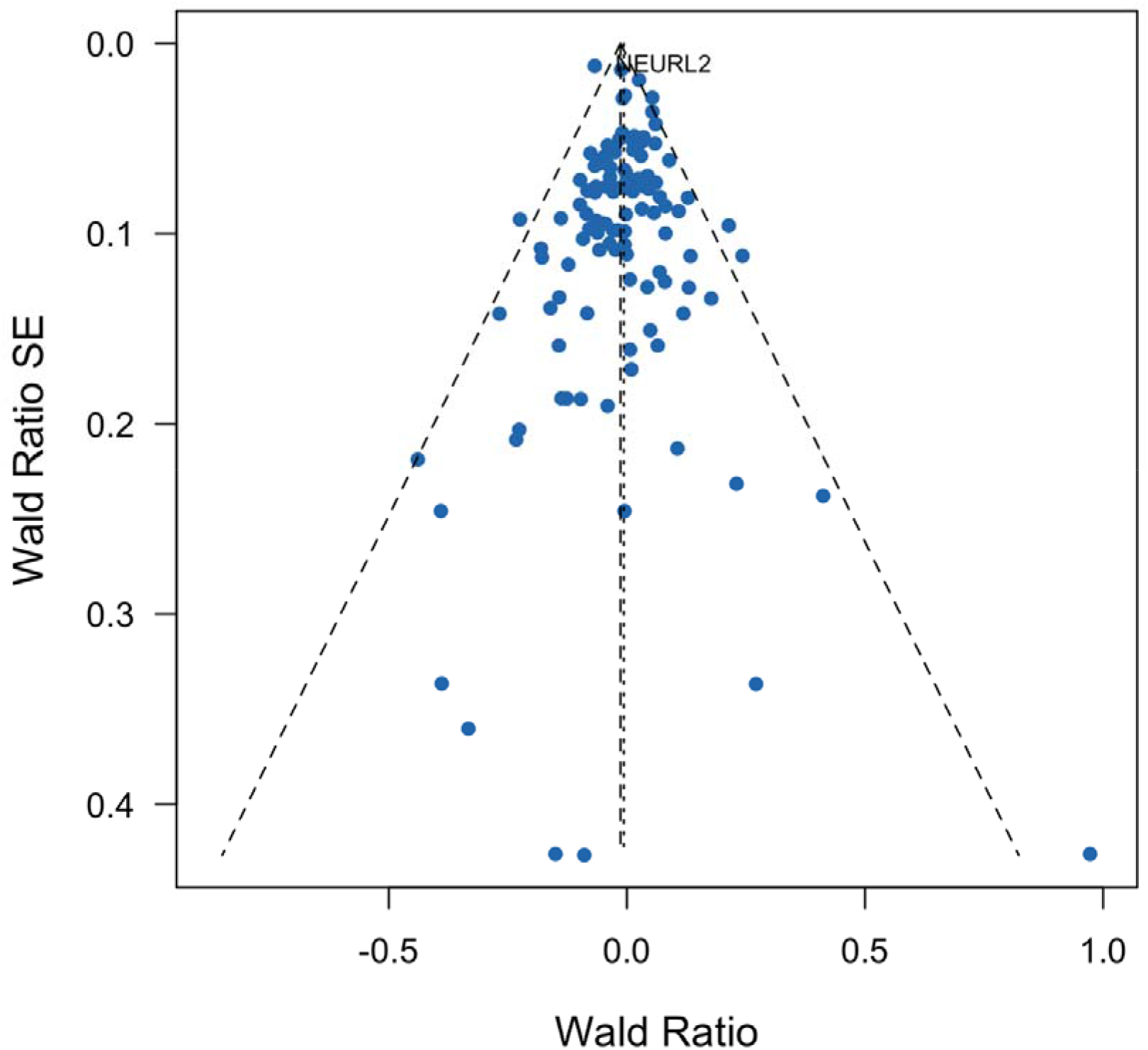
Gene perturbations strongly associated with free-fed glucose in mice and their association with TG:HDL in humans Supplementary Figure 15 presents Wald ratios and standard errors for individual genes strongly associated with free-fed glucose in mice with respect to TG:HDL in humans. Wald ratios are the gene-phenotype effect estimate in humans divided by the gene-phenotype effect estimate in mice. We used directional, unweighted mouse estimates and so the Wald ratios are the effect estimates from humans with positive (consistent directionality between mice and humans) or negative (inconsistent directionality) values. The dotted vertical lines indicate the overall fixed and random effects GPAT estimates across all genes. All genes with pLoF burden P < 0.001 in humans are annotated with the human gene symbol.

